# The host GTPase Dynamin 2 modulates apical junction structure to control cell-to-cell spread of *Listeria monocytogenes*

**DOI:** 10.1101/2024.04.10.588880

**Authors:** Serena Tijoriwalla, Thiloma Liyanage, Thilina U.B. Herath, Nicole Lee, Attika Rehman, Antonella Gianfelice, Keith Ireton

## Abstract

The food-borne pathogen *Listeria monocytogenes* uses actin-based motility to generate plasma membrane protrusions that mediate the spread of bacteria between host cells. In polarized epithelial cells, efficient protrusion formation by *Listeria* requires the secreted bacterial protein InlC, which binds to a carboxyl-terminal Src Homology 3 (SH3) domain in the human scaffolding protein Tuba. This interaction antagonizes Tuba, thereby diminishing cortical tension at the apical junctional complex and enhancing *L. monocytogenes* protrusion formation and spread. Tuba contains five SH3 domains apart from the domain that interacts with InlC. Here we show that the human GTPase Dynamin 2 associates with two SH3 domains in the amino terminus of Tuba and acts together with this scaffolding protein to control spread of *L. monocytogenes*. Genetic or pharmacological inhibition of Dynamin 2 or knockdown of Tuba each restored normal protrusion formation and spread to a bacterial strain deleted for the *inlC* gene (Δ*inlC*). Dynamin 2 localized to apical junctions in uninfected human cells and to protrusions in cells infected with *Listeria*. Localization of Dynamin 2 to junctions and protrusions depended on Tuba. Knockdown of Dynamin 2 or Tuba diminished junctional linearity, indicating a role for these proteins in controlling cortical tension. Collectively, our results show that Dynamin 2 cooperates with Tuba to promote intercellular tension that restricts spread of Δ*inlC Listeria*. By expressing InlC, wild-type *L. monocytogenes* overcomes this restriction.

## INTRODUCTION

*Listeria monocytogenes* is a food-borne pathogen that causes gastroenteritis, meningitis, or abortion (1). This bacterium uses an actin-based motility process to spread between host cells in the intestinal epithelium and liver, and to traverse the placental barrier (2–4). Cell-to-cell spread of *Listeria* is initiated by the microbial surface protein ActA, which stimulates the formation of filamentous (F)-actin tails that propel bacteria through the cytosol of the host cell. Motile bacteria reach the cell periphery, where they remodel the plasma membrane into protrusions that are internalized by adjacent host cells. In this way, actin-based motility and the subsequent formation of membrane protrusions result in intercellular spread of bacteria.

*Listeria* initiates disease by infecting the intestinal epithelium, which consists of polarized cells that are organized into a tight barrier due to the apical junctional complex (AJC) (5–7). The AJC consists of tight junctions (TJs) and underlying adherens junctions, which are linked to an actomyosin network that generates cortical tension (8–12). This tension contributes to the barrier properties of epithelia (8, 10, 13) and has the potential to limit intercellular spread of pathogens by reducing pliability of the plasma membrane. However, work with the polarized human epithelial cell line Caco-2 BBE1 has shown that *Listeria* secretes a virulence protein called InlC that relieves cortical tension, thereby allowing bacteria to spread efficiently (14, 15).

InlC affects tension and promotes *Listeria* protrusion formation by antagonizing the human scaffolding protein Tuba (15, 16). Tuba contains several functional regions, including a Bin-Amphiphysin-Rvs (BAR) domain that binds membranes, a Dbl homology (DH) domain that activates the GTPase Cdc42, and six Src Homology 3 (SH3) domains (17). An SH3 domain located in the carboxyl-terminus of Tuba called “SH36” interacts with the human actin regulatory protein N-WASP, forming a complex that contributes to cortical tension (12, 17). Upon infection with *Listeria*, InlC binds directly to SH36 and displaces N-WASP from Tuba (15, 16). This activity attenuates tension, resulting in enhanced protrusion formation and cell-to-cell spread of *Listeria*.

The amino-terminus of Tuba contains four tandem SH3 domains that directly interact with the mammalian GTPase Dynamin 1 (17). Dynamin 1 is the founding member of a GTPase family that also includes Dynamin 2 and Dynamin 3 (18, 19). Whereas Dynamin 1 and Dynamin 3 are expressed mainly in the brain, expression of Dynamin 2 is ubiquitous. The three Dynamin GTPases each have important roles in clathrin-mediated endocytosis (CME), albeit in different cell types (18–20). Dynamin 1 and Dynamin 3 promote endocytosis and recycling of synaptic vesicles in presynaptic neurons, while Dynamin 2 mediates CME in non-neuronal cells (20–22). Dynamin 2 also contributes to endocytic pathways that are independent of clathrin, including caveolin-mediated endocytosis, micropinocytosis, and internalization of select cytokine receptors (23–25). The three Dynamin proteins exhibit GTP-dependent membrane fission activity, which is critical for their endocytic roles (19). In addition to stimulating membrane fission, Dynamin proteins use their GTPase activity to regulate the actin cytoskeleton (18, 26, 27). This cytoskeletal function contributes to endocytosis (26) and also controls cell migration (28), myoblast formation (27), tight junction structure (29–31), and the generation of protrusive structures such as podosomes (32–35), dendritic spines (36), and axonal growth cones (37, 38).

The three Dynamin proteins share a common structure that includes an amino-terminal domain with GTPase activity, a central ‘stalk’ region containing a four-helix bundle, and a carboxyl-terminal proline and arginine-rich domain (PRD) (19). Dynamin GTPases are recruited to the plasma membrane by interaction of their PRD with SH3 domains in various proteins that contain BAR (Bin-Amphiphysin-Rvs) domains (39, 40). Dynamin proteins then oligomerize, resulting in stimulation of GTP hydrolysis and consequent conformational changes that promote membrane fission or actin polymerization (19, 26, 27).

In this work, we used biochemical, genetic, and microscopy-based approaches to investigate if the ubiquitously expressed GTPase Dynamin 2 acts together with the BAR domain protein Tuba to control intercellular spread of *Listeria* in Caco-2 BBE1 cells. Our results indicate that interaction of the PRD in Dynamin 2 with SH3 domains in the amino-terminus of Tuba recruits Dynamin 2 to the AJC. There, both Dynamin 2 and Tuba contribute to cortical tension that inhibits protrusion formation and spread of a *Listeria* mutant strain lacking the virulence gene *inlC*. Wild-type bacteria that produce InlC overcome this inhibition and undergo efficient intercellular spread.

## RESULTS

### SH3 domains in the amino terminus of Tuba associate with Dynamin 2

In humans and mice, Tuba is expressed as two different isoforms derived from alternative splicing (17). The largest isoform, referred to here as isoform 1, has an amino-terminal region containing four tandem SH3 domains (SH31-4), a Dbl Homology (DH) domain that activates the GTPase Cdc42, a BAR domain, and two SH3 domains (SH35 and SH36) located in the carboxyl-terminal end of the protein (Fig. 1A). Previous findings indicate that the SH31-4 region binds to Dynamin 1 (17), a GTPase expressed mainly in neuronal cells (19). In order to better understand how Tuba controls cell-to-cell spread of *Listeria*, we investigated if this same region in Tuba associates with Dynamin 2, a ubiquitously expressed protein that is closely related to Dynamin 1 (19).

**Figure 1.**
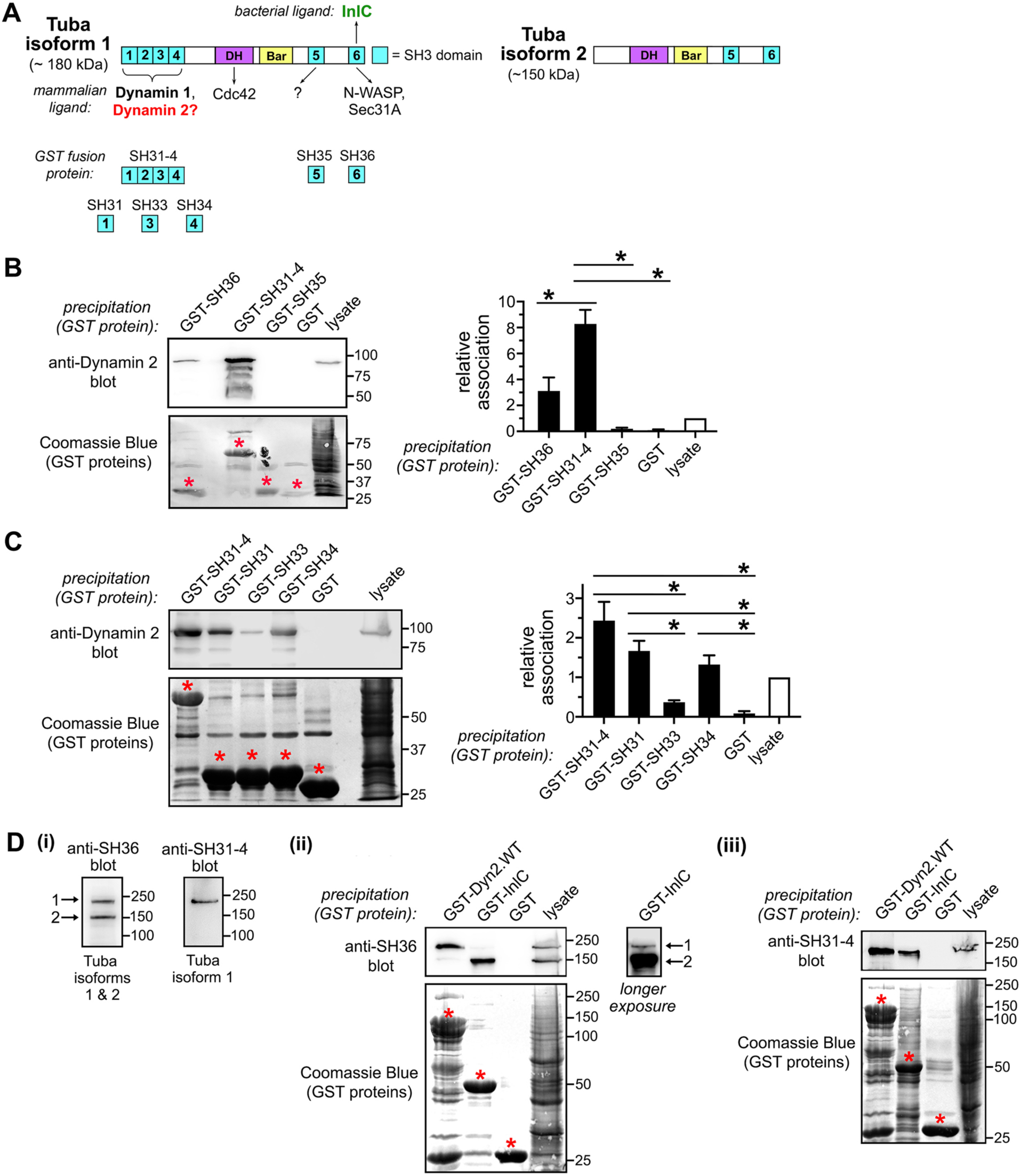
Dynamin 2 associates with SH3 domains in the amino terminus of Tuba. A. Structures of Tuba proteins expressed in Caco-2 BBE1 cells. The two Tuba isoforms are distinguished by the presence or absence of the SH31-4 region (15, 17, 67). Mammalian and bacterial ligands that interact with domains in Tuba are depicted. GST fusion proteins with SH3 domains of Tuba used for co-precipitation studies are shown below Tuba isoform 1. B. Co-precipitation of human Dynamin 2 with the SH31-4, SH35, or SH36 regions in Tuba. Lysates of Caco-2 BBE1 cells were used for precipitation with GST alone or GST fused to the indicated domains in Tuba, followed by Western blotting with antibodies recognizing Dynamin 2. To confirm precipitation of GST proteins, membranes were stripped and stained with Coomassie Blue. Panels on the left are representative results. Red asterisks indicate precipitated GST fusion proteins. The graph on the right has co-precipitation data from four experiments. *, P < 0.05. C. The ability of individual SH3 domains in the SH31-4 region to precipitate Dynamin 2 from Caco-2 BBE1 cell lysates. Data in the graph are means +/- SEM from three experiments. *, P < 0.05. D. Selective association of Dynamin 2 with isoform 1 of Tuba. (i). Western blots of Caco-2 BBE1 cell lysates using antibodies raised against the SH31-4 or SH36 regions in Tuba. Anti-SH36 antibodies detect both Tuba isoforms, whereas anti-SH31-4 antibodies recognize only Tuba isoform 1 (15) (ii). Caco-2 BBE1 cell lysates were used for precipitation with the indicated GST protein, followed by Western blotting of precipitates with anti-SH36 antibodies. (iii) GST protein precipitates were immunoblotted with anti-SH31-4 antibodies. Red asterisks indicate precipitated GST proteins. In the case of GST-Dyn2.WT, the upper band represents the full-length protein, whereas the smaller bands are likely degradation products. The data in ii or iii are representative of three or four experiments, respectively.

Co-precipitation experiments were performed using proteins consisting of an amino-terminal glutathione S-transferase (GST) tag fused to the SH31-4, SH35, or SH36 domains in Tuba (Fig. S1A). Lysates of Caco-2 BBE1 cells were incubated with these proteins or GST alone as a control, followed by precipitation of GST proteins and detection of Dynamin 2 by Western blotting (Fig. 1B). The results indicate that Dynamin 2 associated with the SH31-4 region of Tuba and, to a lesser extent, with the SH36 domain. Individual SH3 domains in the SH31-4 region were then assessed for interaction with Dynamin 2. GST-tagged proteins containing the first, third, and fourth SH3 domains (SH31, SH33 and SH34) were purified (Fig. S1B). A GST fusion protein with the second domain (SH32) could not be purified, possibly because of accumulation in inclusion bodies in *E. coli*. Results from co-precipitation experiments showed that Dynamin 2 associated mainly with the SH31 and SH34 domains (Fig. 1C).

Given that isoform 2 of Tuba lacks the amino-terminal SH31-4 region (Fig. 1A), we anticipated that this isoform would not interact efficiently with Dynamin 2. To test this idea, co-precipitation experiments were performed using GST-tagged wild-type (WT) Dynamin 2 protein (Fig. S1C) and antibodies that recognize the SH36 or SH31-4 regions in Tuba (15). The anti-SH36 antibodies detect both Tuba isoforms, whereas the anti-SH31-4 antibodies recognize only Tuba isoform 1 (Fig. 1D part i). As a positive control, a GST fusion protein containing the *Listeria* virulence protein InlC was included. This GST-InlC protein binds to the SH36 domain present in both Tuba isoforms (15, 16). Results from the co-precipitation experiments indicated that GST-Dynamin 2.WT associates exclusively with Tuba isoform 1 (Fig. 1D parts ii and iii).

Dynamin 2 comprises an amino-terminal GTPase (G) domain, a middle region, a pleckstrin homology (PH) domain, a GTPase effector domain (GED), and a carboxyl terminal proline arginine -rich domain (PRD) (19) (Fig. 2A). The PRD contains several peptides of the sequences RxxPxxP or xPxxPxR (where x is any amino acid) that bind SH3 domains in proteins involved in endocytosis and/or regulation of the actin cytoskeleton (26). To examine if the PRD in Dynamin 2 mediates interaction with SH3 domains in Tuba, the human epithelial cell line HeLa was transfected with plasmids expressing proteins consisting of wild-type (WT) Dynamin 2 or Dynamin 2 lacking the PRD (ΔPRD) fused to enhanced green fluorescent protein (EGFP) (Fig. 2A). Western blotting of total cell lysates indicated that the WT and ΔPRD Dynamin 2 proteins were expressed at similar levels (Fig. 2B). Co-precipitation experiments with GST fusion proteins showed that the SH31-4 region, SH31 domain, or SH34 domain associated more efficiently with WT Dynamin 2-EGFP compared to Dynamin 2ΔPRD-EGFP (Fig. 2C). In addition, Tuba isoform 1 from Caco-2 BBE1 cell lysates co-precipitated to a greater extent with GST-tagged WT Dynamin 2 protein than with GST-Dynamin 2ΔPRD (Fig. 2D). Collectively, the data in Figure 2 indicate that the PRD in Dynamin 2 is needed for efficient interaction with Tuba.

**Figure 2.**
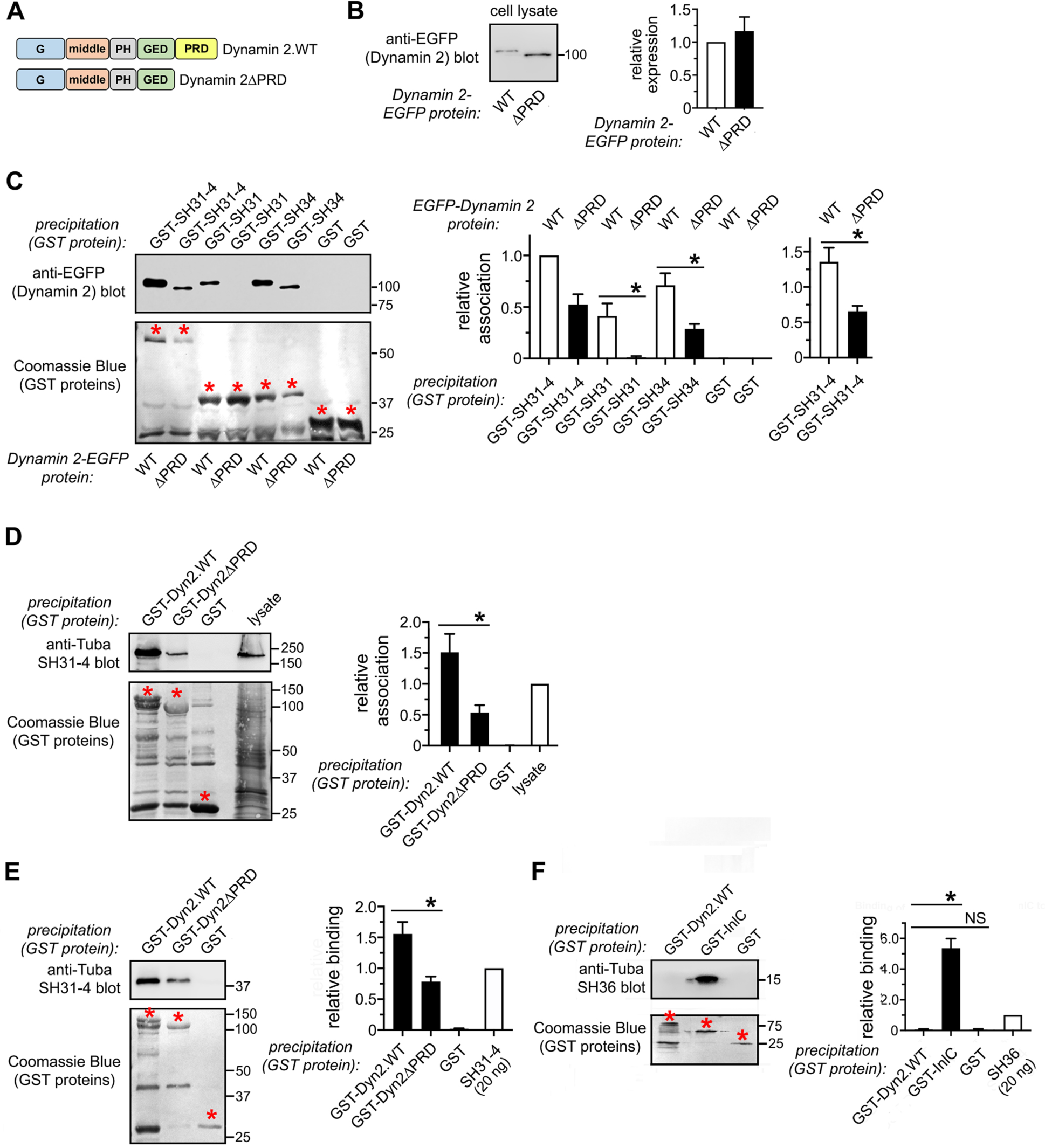
The PRD in Dynamin 2 contributes to interaction with Tuba. A. Structures of wild-type (WT) Dynamin 2 and Dynamin 2 deleted for its PRD (ΔPRD) used in co-precipitation studies. B. Expression of EGFP-tagged Dynamin 2.WT or EGFP-Dynamin 2ΔPRD in transiently transfected HeLa cells is shown. The left panel is a representative image, and the graph contains quantified Western blotting data from five experiments. C. Role of the PRD in interaction with the Tuba SH31-4 region. Lysates from HeLa cells transfected with the EGFP-tagged Dynamin 2 constructs in part B were used for co-precipitation with GST alone or GST proteins containing the indicated domains in Tuba. EGFP-tagged Dynamin 2 proteins were detected by Western blotting. The left panels are representative results, and the graph contains mean +/- SEM values from four experiments. *, P < 0.05. Red asterisks indicate precipitated GST fusion proteins. D. Contribution of the PRD in association with Tuba isoform 1. Lysates of Caco-2 BBE1 cells were used for precipitation with the indicated GST fusion protein. The presence of Tuba isoform 1 in precipitates was detected by Western blotting with antibodies against the Tuba SH31-4 region. Data in the graph are mean +/- SEM values from four experiments. *, P < 0.05. E. Direct binding of Dynamin 2 to the Tuba SH31-4 region. Approximately 30 nM of GST-Dynamin 2.WT, GST-Dynamin 2ΔPRD, or GST alone were incubated with 3 nM of the SH31-4 region of Tuba for 2 hours, followed by precipitation of GST fusion proteins and detection of SH31-4 by Western blotting. (i) A representative Western blot image is shown. (ii) The graph contains mean +/- SEM values of co-precipitation of SH31-4 from four experiments. *, P <0.05. F. Lack of binding of Dynamin 2 to the Tuba SH36 domain. Approximately 24 nM of the SH36 domain of Tuba was incubated with 30 nM of GST-Dynamin 2.WT, GST-InlC, or GST alone, followed by precipitation and Western blotting to detect SH36. The graph shows results from three experiments. *, P <0.05. NS indicates not significant.

### Interaction between Dynamin 2 and the Tuba SH31-4 region is direct

In order to determine if the SH31-4 region interacts directly with Dynamin 2, co-precipitation assays were performed using purified GST-Dynamin 2.WT or GST-Dynamin 2ΔPRD and SH31-4 with its GST tag removed (Fig. S1C,D). The results showed that SH31-4 co-precipitated with GST-Dynamin 2.WT, but not with GST alone (Fig. 2E). Co-precipitation of SH31-4 with GST-Dynamin 2.WT was more efficient than with GST-Dynamin 2 ΔPRD. We conclude that Dynamin 2 binds directly to SH31-4 in a manner partly dependent on the PRD.

Since GST fused to the SH36 domain of Tuba co-precipitated weakly with Dynamin 2 from lysates of Caco-2 BBE1 cells (Fig. 1B), we used purified SH36 with its GST tag removed (Fig. S1D) to determine if the SH36 region binds directly to purified GST-Dynamin 2.WT. As a positive control, GST fused to InlC was used. Whereas SH36 interacted with GST-InlC, this SH3 domain failed to co-precipitate with GST-Dynamin 2.WT (Fig. 2F). The weak association between GST-SH36 and Dynamin 2 in cell lysates (Fig. 2B) therefore likely reflects an indirect interaction, which presumably involves an intermediary protein. The identity of such a protein is presently unknown.

### Dynamin 2 and Tuba have similar roles in protrusion formation and cell-to-cell spread of *Listeria*

Results from a previous study indicated that Tuba antagonizes protrusion formation and cell-to-cell spread of a *Listeria* mutant strain that lacks the bacterial virulence gene *inlC* (15). Specifically, it was found that, compared to wild-type *Listeria*, an isogenic mutant strain deleted for *inlC* (Δ*inlC*) exhibits a 40-50% reduction in protrusion formation and spread in Caco-2 BBE1 cells. However, depletion of Tuba by RNAi restores the production of protrusions by the Δ*inlC* mutant to normal levels. We used RNAi to assess if Dynamin 2 and Tuba control protrusion formation and spread in a similar manner. In order to control for potential off-target effects of short interfering RNAs (siRNAs) two different siRNAs targeting distinct regions of Dynamin 2 mRNA called Dyn2-2 and Dyn2-3 were used. As controls, cells were mock transfected in the absence of siRNA or transfected with a control, non-targeting siRNA which lacks complementarity to any known human mRNA. Protrusion formation efficiency was assessed by determining the percentage of bacterial-associated F-actin structures in protrusions, as previously described (Fig. S2) (15, 16, 41–43). Cell-to-cell spread was quantified using focus assays, which relate the efficiency of bacterial spread in a cell monolayer to the size infection foci (Fig. S3) (15, 16, 41–43). The results indicated that siRNA-mediated depletion of Dynamin 2 or Tuba cause similar restorations in the formation of protrusions and intercellular spread of the Δ*inlC* strain of *Listeria* (Fig. 3A,B and Fig. 4). Importantly, depletion of Dynamin 2 or Tuba did not alter the frequency of bacteria that recruit F-actin in the cytosol or the mean lengths of actin comet tails in the main body of the host cell (Fig. S4 parts A, B, and C). These results show that knockdown of Dynamin 2 or Tuba did not affect escape of bacteria to the cytosol or the rate of actin-based motility, which is proportional to comet tail lengths (44). We therefore conclude that Dynamin 2 and Tuba directly control protrusion formation by the Δ*inlC* strain of *Listeria*.

**Figure 3.**
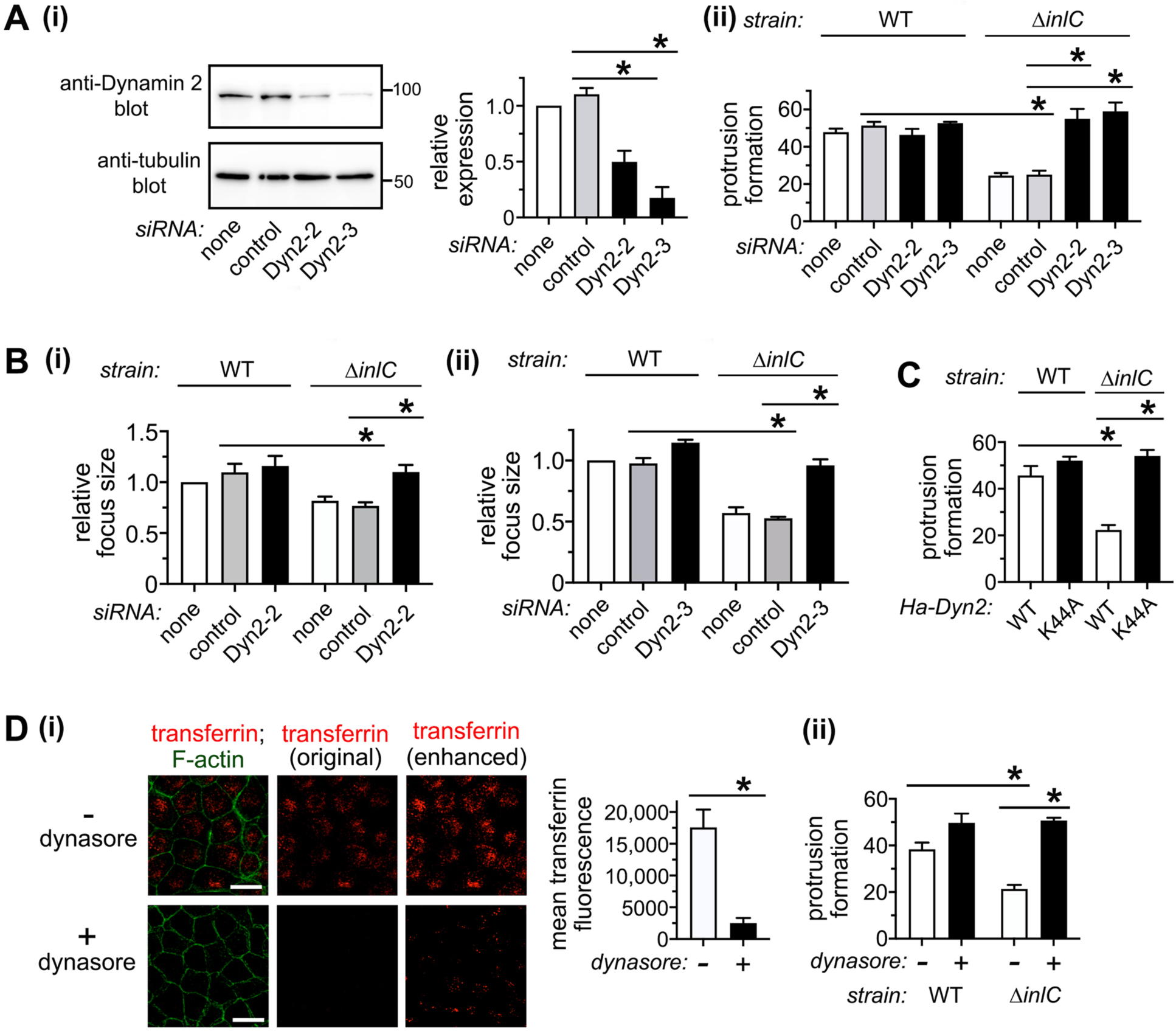
Dynamin 2 restricts intercellular spread of bacteria lacking InlC. A. Effect of siRNAs targeting Dynamin 2 on target protein expression (i) and protrusion formation by *L. monocytogenes* (ii). Caco-2 BBE1 cells were either left untransfected (none), transfected with a control non-targeting siRNA, or transfected with two different siRNAs against Dynamin 2. Approximately 72 h later, cells were solubilized in lysis buffer for assessment of depletion of Dynamin 2 by Western blotting (i) or infected with wild-type (WT) or Δ*inlC* strains of *L. monocytogenes* for 6 hours for measurement of protrusion formation efficiency, as described (15, 16, 41, 43) (ii). The Western blotting and protrusion formation results are mean +/- SEM values from three experiments. B. Effect of two different siRNAs against Dynamin 2 on cell-to-cell spread of *L. monocytogenes* strains expressing or lacking InlC. Caco-2 BBE1 cells transfected with Dyn2-2 siRNA (i) or Dyn2-3 siRNA (ii) or subjected to control conditions were infected with WT or Δ*inlC* bacterial strains for 12 h for measurement of cell-to-cell spread using focus assays (16, 41, 43). Data are mean focus size -/+ SEM from three experiments. In each experiment, approximately 25 foci were measured for each condition. C. Impact of a Dynamin 2 protein defective in GTP binding on *L. monocytogenes* protrusion formation. Caco-2 BBE1 cells were transfected with plasmids expressing Ha epitope-tagged wild-type (WT) Dynamin 2 or Dynamin 2.K44A, followed by infection with the indicated *L. monocytogenes* strains. Data are mean -/+ SEM values from three experiments. D. Effect of dynasore treatment on bacterial protrusions. Caco-2 BBE1 cells were incubated in medium containing 0.2% DMSO (-) or 80 µM dynasore for 4.5 hours, followed by assessment of endocytosis of transferrin (i) or protrusion formation by WT or Δ*inlC* strains of *L. monocytogenes* (ii). For quantification of internalized transferrin, original (non-enhanced) images were used. A digitally enhanced image is included to facilitate visualization of labeled transferrin. Data in i and ii are mean +/- SEM values from three experiments. Scale bars are 20 micrometers. *, P < 0.05.

**Figure 4.**
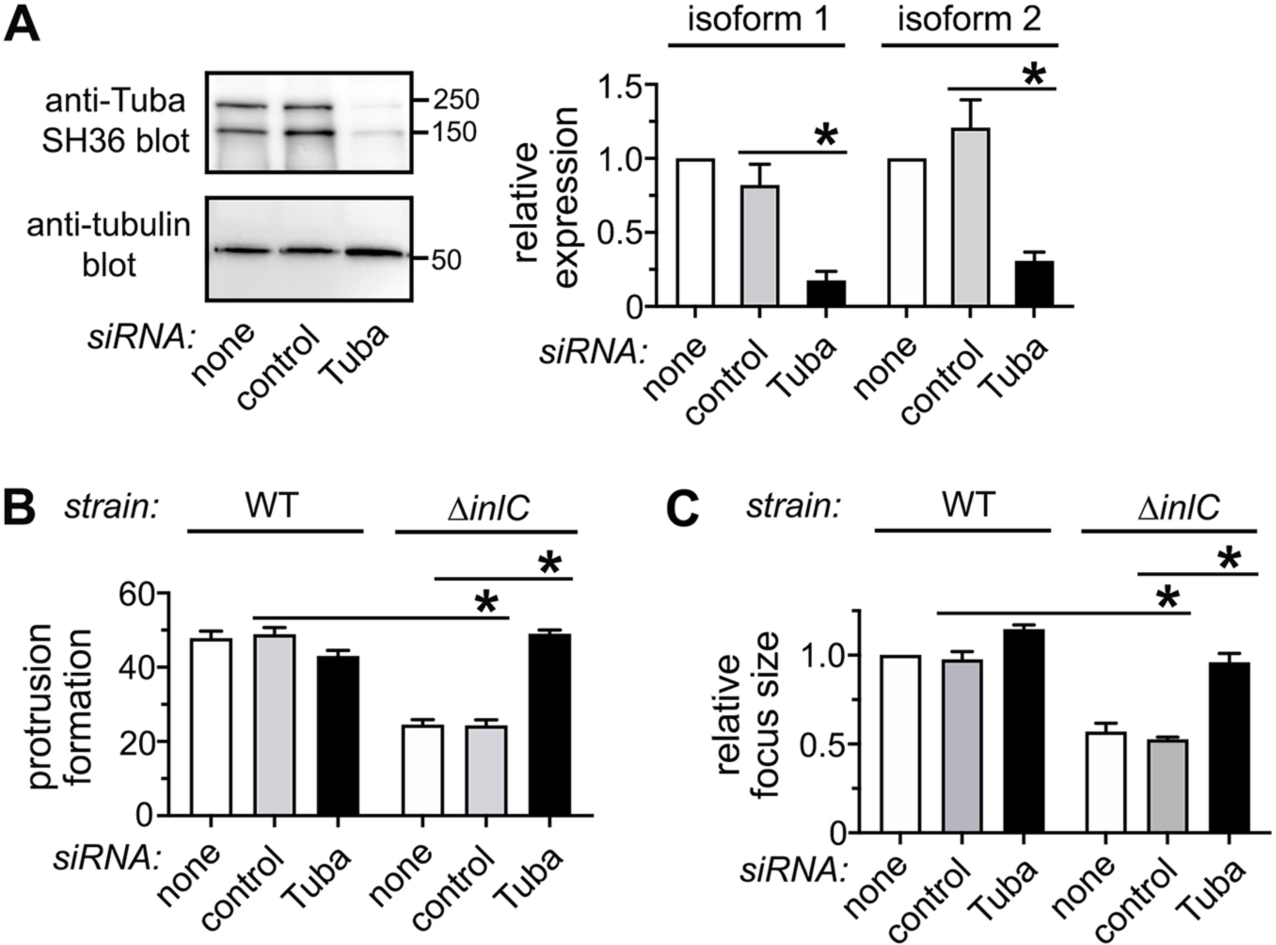
Tuba controls *L. monocytogenes* spread similarly to Dynamin 2. Caco-2 BBE1 cells were either left untransfected (none), transfected with a control siRNA, or transfected with an siRNA targeting Tuba for about 72 h. Cells were then processed for assessment of Tuba protein expression through Western blotting (A), protrusion formation by WT and Δ*inlC* strains of *L. monocytogenes* (B), or cell-to-cell spread of bacteria (C). The data are means +/- SEM from three experiments. In C, approximately 25 foci were analyzed per condition in each experiment. *, P < 0.05.

As previously reported (15), depletion of Tuba did not impact protrusion formation or spread by the WT *Listeria* strain (Fig. 4). RNAi-induced knockdown of Dynamin 2 also failed to affect infection by this strain (Fig. 3A,B). These results are consistent with previous biochemical and genetic data showing that InlC protein produced by WT *Listeria* binds directly to Tuba and antagonizes its function (15, 16). This antagonism explains why knockdown of Tuba or Dynamin 2 does not alter protrusion formation or spread of WT bacteria.

We next examined if the ability of Dynamin 2 to bind and hydrolyze GTP impacts protrusion formation by *Listeria*. For these studies, we used a commonly employed mutant Dynamin 2 protein defective in GTP binding (Dynamin 2.K44A) or a chemical inhibitor of Dynamin protein GTPase activity called dynasore (26, 45, 46). Transient expression of Dynamin 2.K44A containing an amino-terminal Ha tag restored normal protrusion formation to the Δ*inlC Listeria* strain (Fig. 3C), without affecting escape to the cytosol or actin tail length (Fig. S4D). Treatment of Caco-2 BBE1 cells with dynasore caused similar effects on production of protrusions (Figs. 3D and S4E). In these latter experiments, the activity of dynasore was confirmed by demonstrating inhibition of endocytosis of transferrin at the same concentration used in the infection studies. Taken together, the findings in Figures 3 and 4 demonstrate that Dynamin 2 and Tuba control *Listeria* protrusion formation and cell-to-cell spread in a similar manner. The results also suggest that regulation of infection by Dynamin 2 requires its ability to bind and hydrolyze GTP.

### Dynamin 2 localizes to protrusions in cells donating these structures

We examined localization of Dynamin 2 during infection with WT or Δ*inlC* strains of *Listeria*. Caco-2 BBE1 cells were transfected with plasmids expressing EGFP-tagged WT Dynamin 2 or Dynamin 2ΔPRD and then infected with WT or Δ*inlC* strains of *Listeria* for 6 h, followed by processing of samples for laser scanning confocal microscopy analysis. In resulting images, protrusions were identified as EGFP-positive structures projecting from human cells expressing EGFP into adjacent host cells lacking EGFP. Dynamin 2.WT-EGFP was observed to accumulate extensively in protrusions of WT *Listeria* (Fig. 5A). The degree of accumulation of WT Dynamin 2 or Dynamin 2ΔPRD in protrusions was quantified as fold enrichment (FE) values. This method has been previously used to quantify recruitment of other human proteins to membrane protrusions generated by *L. monocytogenes* or the bacterium *Shigella flexneri* (41, 47). FE is defined as the ratio of the mean pixel intensity (mpi) of a protein of interest in protrusions divided by the mpi of the protein throughout the entire host cell. An FE value greater than 1.0 indicates enrichment of the protein in protrusions. Quantification of FE values for ∼ 30 protrusions indicated that Dynamin 2.WT-EGFP had average enrichment of about two-fold in protrusions of WT *Listeria*, whereas Dynamin 2ΔPRD-EGFP was not enriched (Fig. 5B). These data indicate that the PRD mediates recruitment of Dynamin 2 to protrusions. A comparison of FE values for protrusions made by WT or Δ*inlC* strains of *Listeria* revealed that InlC contributes to the recruitment of WT Dynamin 2 to protrusions (Fig. 5B). We also examined localization of endogenous Dynamin 2 protein by labeling cells with an antibody against the GTPase (Fig. S5). In these studies, protrusions were identified by transfecting Caco-2 BBE1 cells with a plasmid expressing EGFP and assessing Dynamin 2 labeling in projections emanating from EGFP-positive cells into surrounding cells. In agreement with the data with Dynamin 2.WT-EGFP, the results show that endogenous Dynamin 2 is enriched to a greater extent in protrusions of WT *Listeria* compared to those of Δ*inlC* bacteria. Collectively, the findings in Figure 5 and S5 indicate important roles for the PRD and InlC in mobilization of Dynamin 2 during cell-to-cell spread.

**Figure 5.**
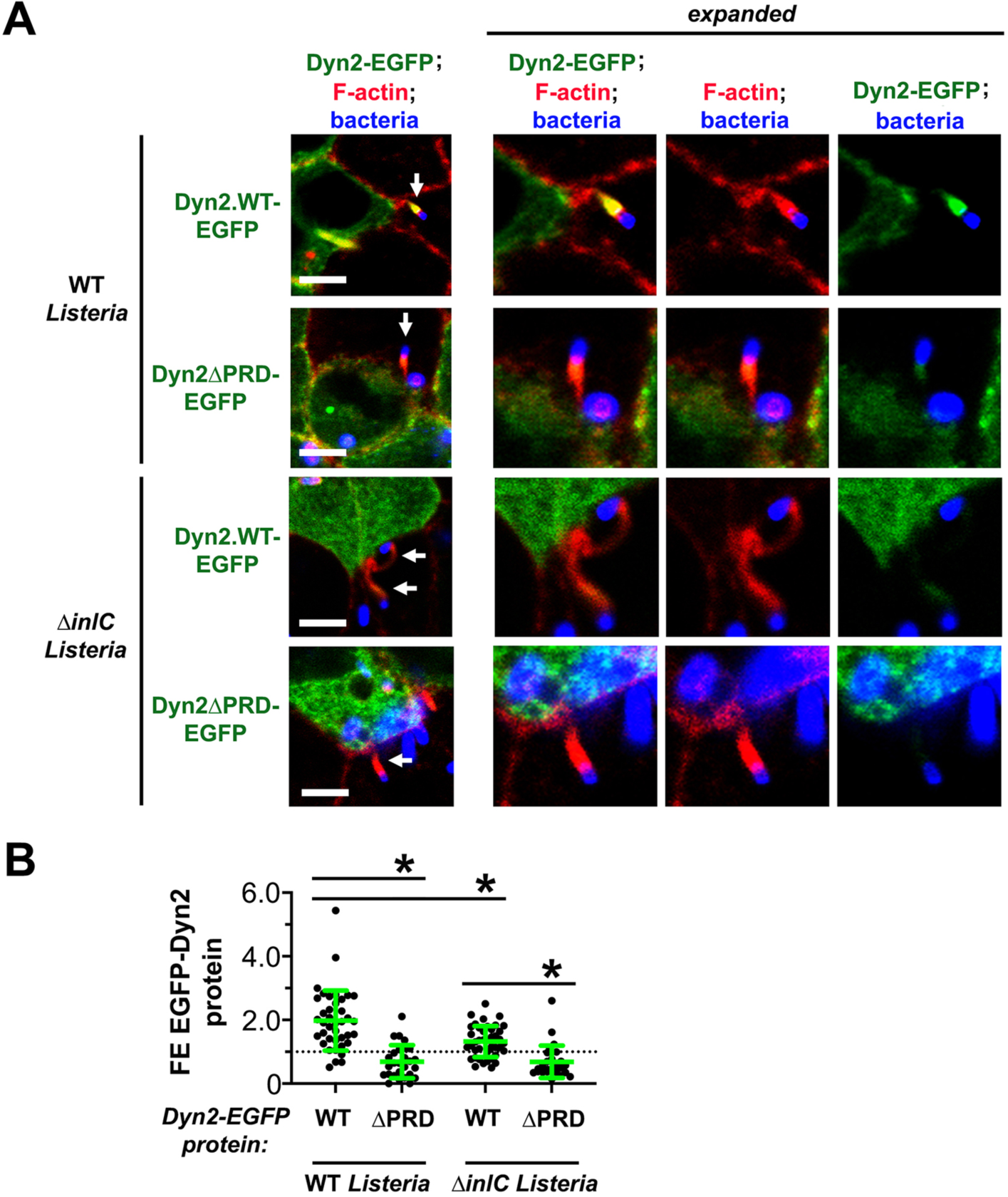
Dynamin 2 is enriched in *L. monocytogenes* protrusions in a manner dependent on the PRD and the bacterial protein InlC. Caco-2 BBE1 cells transiently expressing EGFP-tagged wild-type (WT) Dynamin 2 or Dynamin 2 deleted for its PRD (ΔPRD) were infected with WT or Δ*inlC* strains of *L. monocytogenes* for 6 h. Cells were then fixed, labeled, and imaged using laser scanning confocal microscopy. A. Representative microscopy images are shown. Arrows indicate protrusions projecting from host cells expressing EGFP-tagged Dynamin 2 proteins into surrounding cells lacking these proteins. Panels on the right are expanded to facilitate visualization of protrusions. Scale bars indicate 10 micrometers. B. Enrichment of tagged Dynamin2 proteins in protrusions with *L. monocytogenes*. Each dot is a fold enrichment (FE) value of a protrusion. Between 27-41 FE measurements were made, depending on the condition. Data are means +/- SD. *, P < 0.05.

We considered the possibility that Dynamin 2 might also accumulate around *Listeria* protrusions in Caco-2 BBE1 cells accepting these structures. Such localization of Dynamin 2 was observed in HeLa cells that were in the process of internalizing *Listeria* protrusions (48). We identified images in which protrusions of WT *Listeria* were projecting from a Caco-2 BBE1 cell lacking Dynamin 2-EGFP into a neighboring cell expressing the tagged protein. As a control, images of protrusions projecting from cells expressing Dynamin 2.WT-EGFP into cells lacking this protein were used. The plot profile function of ImageJ software was used to assess the spatial overlap of Dynamin 2.WT-EGFP and F-actin in protrusions. The results indicated that Dynamin 2.WT-EGFP coincided spatially with F-actin in protrusions in host cells donating these structures (Fig. S6A), but not in cells accepting the structures (Fig. S6B). Plot profiles from 25 donated or accepted protrusions were used to determine ratios of Dynamin 2.WT fluorescence intensity in regions overlapping actin-rich protrusions relative to regions outside of protrusions (see Materials and Methods). On average, Dynamin 2.WT-EGFP levels were increased ∼ 3-fold around donated protrusions, but were not augmented around accepted protrusions (Fig. S6C). Collectively, the data in Figures 5 and S6 show that Dynamin 2 accumulates in F-actin tails exclusively in Caco-2 BBE1 cells producing protrusions, which is in general agreement with data indicating that Dynamin 2 controls protrusion formation (Fig. 3).

### Dynamin 2 localizes to F-actin comet tails in the body of infected host cells

In addition to being recruited to protrusions, Dynamin 2.WT-EGFP localized to F-actin tails in the main body of Caco-2 BBE1 cells (Fig. S7), as previously reported for HeLa cells infected with *Listeria* (49, 50). We also detected Dynamin 2.WT-EGFP around bacteria with symmetrically distributed actin, which represents a step preceding actin comet tail formation. Similarly to protrusions, Dynamin 2ΔPRD failed to localize to comet tails or around bacteria with symmetric actin, indicating a critical role for the PRD in recruitment. However, unlike protrusions, the Δ*inlC* mutant strain of *Listeria* did not exhibit reduced accumulation of Dynamin 2.WT-EGFP to comet tails or bacteria with symmetric actin (Fig. S7). These results indicate that the PRD, but not InlC, promotes recruitment of Dynamin 2 during actin-based motility in the cytosol.

### Tuba recruits Dynamin 2 to protrusions

Given that Tuba interacts with Dynamin 2 (Figs. 1 and 2) and that several BAR domain proteins are known to mobilize Dynamin proteins to the plasma membrane (40), we investigated if Tuba mediates the recruitment of Dynamin 2 to *Listeria* protrusions and/or structures associated with actin-based motility in the cytosol. Our results indicate that transfection of Caco-2 BBE1 cells with an siRNA that depletes Tuba reduced accumulation of Dynamin 2.WT-EGFP in protrusions of WT *Listeria* (Fig. 6). By contrast, the same Tuba siRNA failed to affect recruitment of Dynamin 2.WT-EGFP to F-actin comet tails or bacteria decorated with symmetric actin in the main body of host cells (Fig. S8). These findings show that Tuba selectively mobilizes Dynamin 2 to protrusions during cell-to-cell spread.

**Figure 6.**
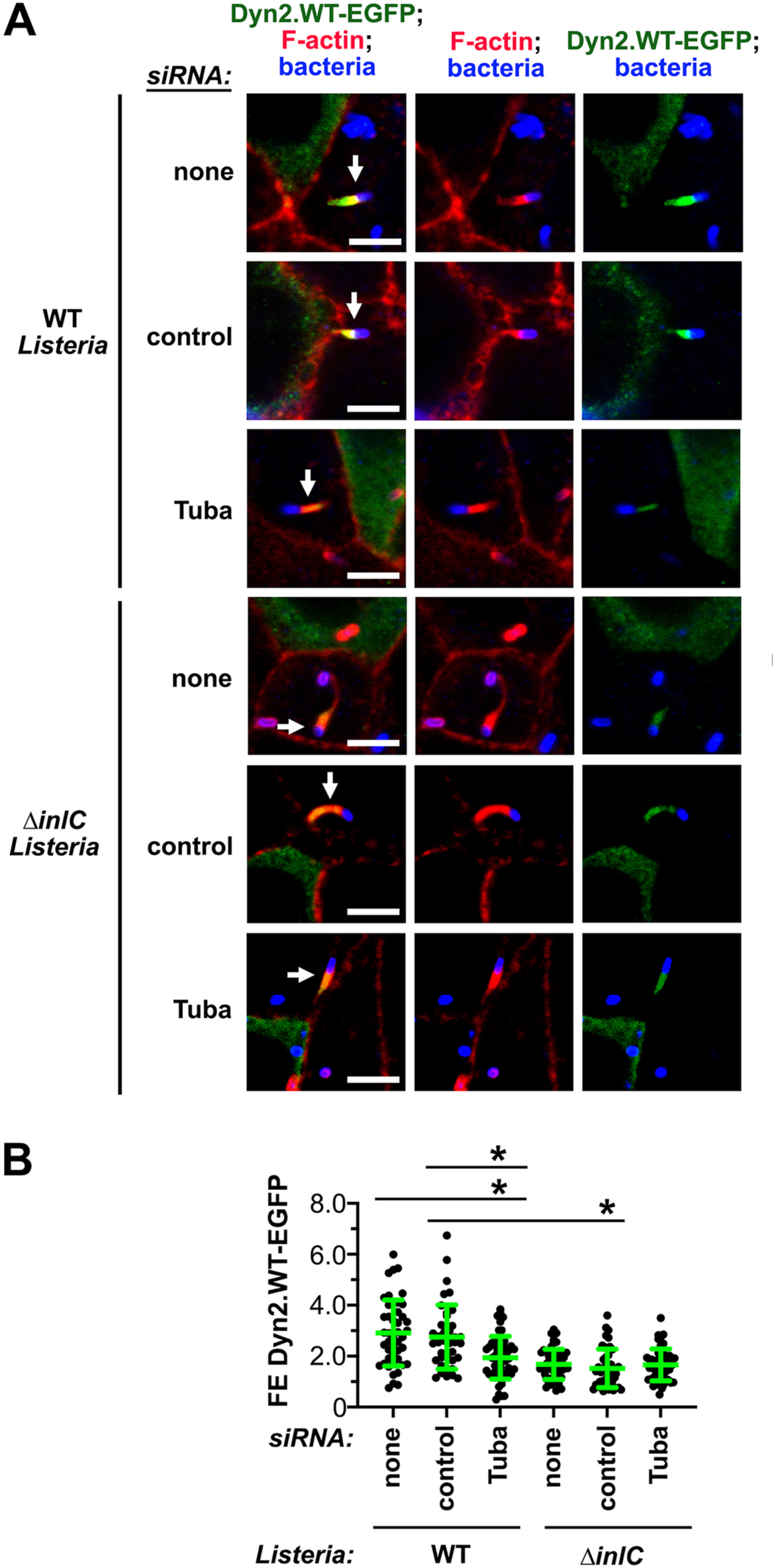
Tuba is needed for recruitment of Dynamin 2 to *L. monocytogenes* protrusions. Caco-2 BBE1 transiently expressing Dynamin 2.WT-EGFP were subjected to control conditions or transfected with an siRNA targeting Tuba. Images of host cells with protrusions labeled with Dynamin 2 were used for quantification of fold-enrichment values, as described (41). A. Representative confocal microscopy images. Protrusions are indicated with arrows. Scale bars represent 10 micrometers. B. The graph contains mean fold enrichment values +/- SEM of 40-55 protrusions. *, P < 0.05.

### Tuba mediates recruitment of Dynamin 2 to tight junctions, where the GTPase controls cortical tension

Tuba localizes to tight junctions of the AJC, where it promotes cortical tension that restrains protrusion formation and intercellular spread of the Δ*inlC* strain of *Listeria* (12, 15, 16, 42, 43). To determine if Dynamin 2 acts together with Tuba at tight junctions to control *Listeria* spread, we investigated if Tuba mobilizes Dynamin 2 to these junctions. Confocal microscopy imaging indicated that Dynamin 2.WT-EGFP, endogenous Dynamin 2, and endogenous Tuba each co-localized with the tight junction protein occludin in Caco-2 BBE1 cells (Fig. 7A,B). Furthermore, endogenous Tuba co-localized with Dynamin 2.WT-EGFP at junctions (Fig. 7C). By contrast, Dynamin 2ΔPRD failed to accumulate at tight junctions or co-localize with Tuba (Fig. 7A,C), indicating a role for the PRD in localization to junctions. Importantly, transfection of cells with an siRNA against Tuba reduced Dynamin 2.WT-EGFP levels at tight junctions (Fig. 8). Collectively, the results in Figures 7 and 8 show that Tuba mediates recruitment of Dynamin 2 to the AJC, probably by interacting with the PRD.

**Figure 7.**
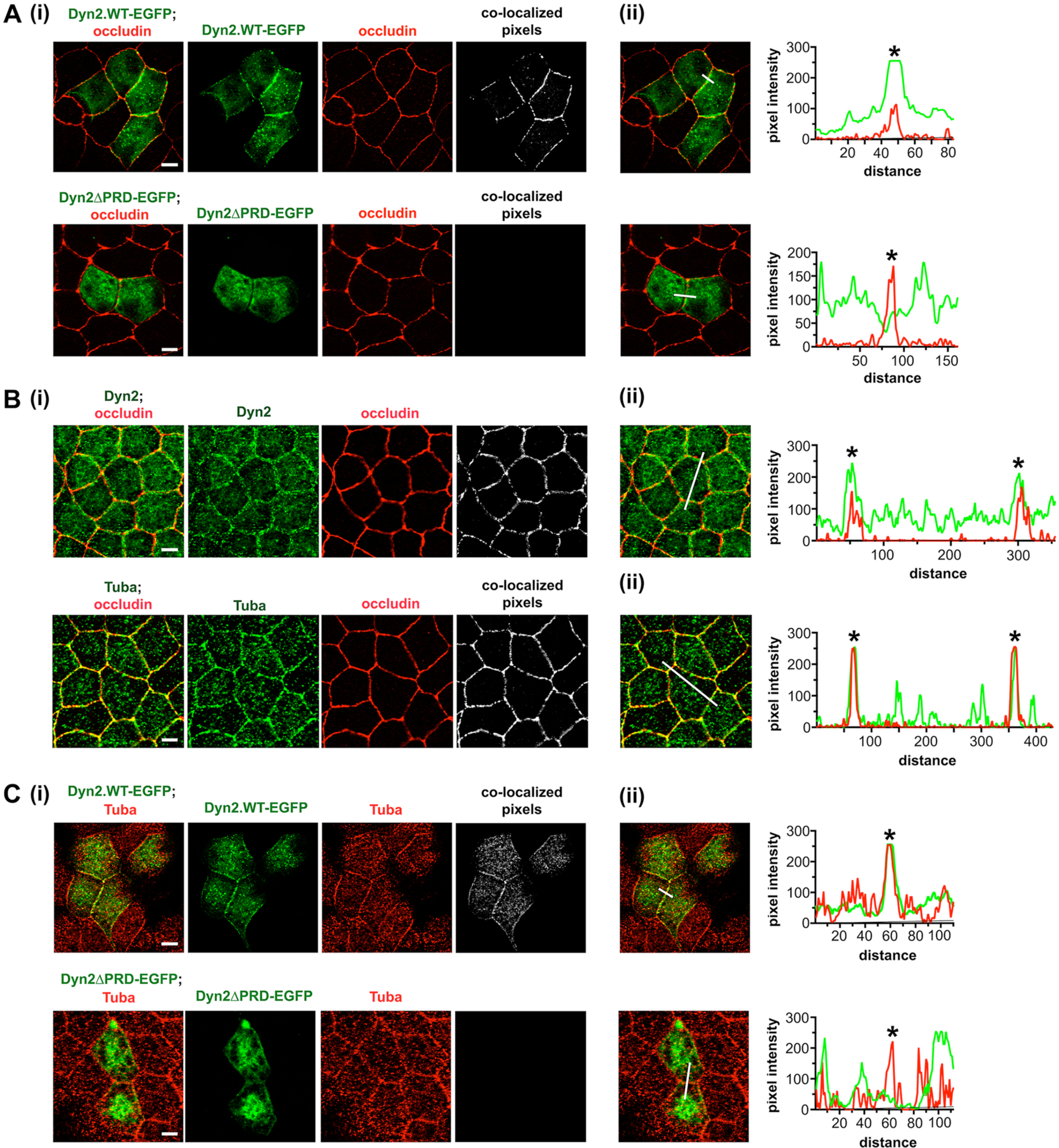
Tuba and Dynamin 2 localize to tight junctions in uninfected cells. A. Co-localization of EGFP-tagged Dynamin 2.WT, but not Dynamin 2ΔPRD, with the tight junction protein occludin. (i) Representative images displaying Dynamin 2-EGFP proteins in green and occludin in red. Co-localized pixels were detected using the co-localization threshold function of ImageJ software. (ii). The plot profile function of ImageJ software was used to display the pixel intensities for EGFP-Dynamin 2 proteins or occludin across the white lines. Asterisks indicate locations of cell junctions. B. Co-localization of endogenous Dynamin 2 or Tuba with occludin. C. Co-localization of Dynamin 2.WT-EGFP, but not Dynamin 2ΔPRD, with endogenous Tuba. All scale bars are 10 micrometers.

**Figure 8.**
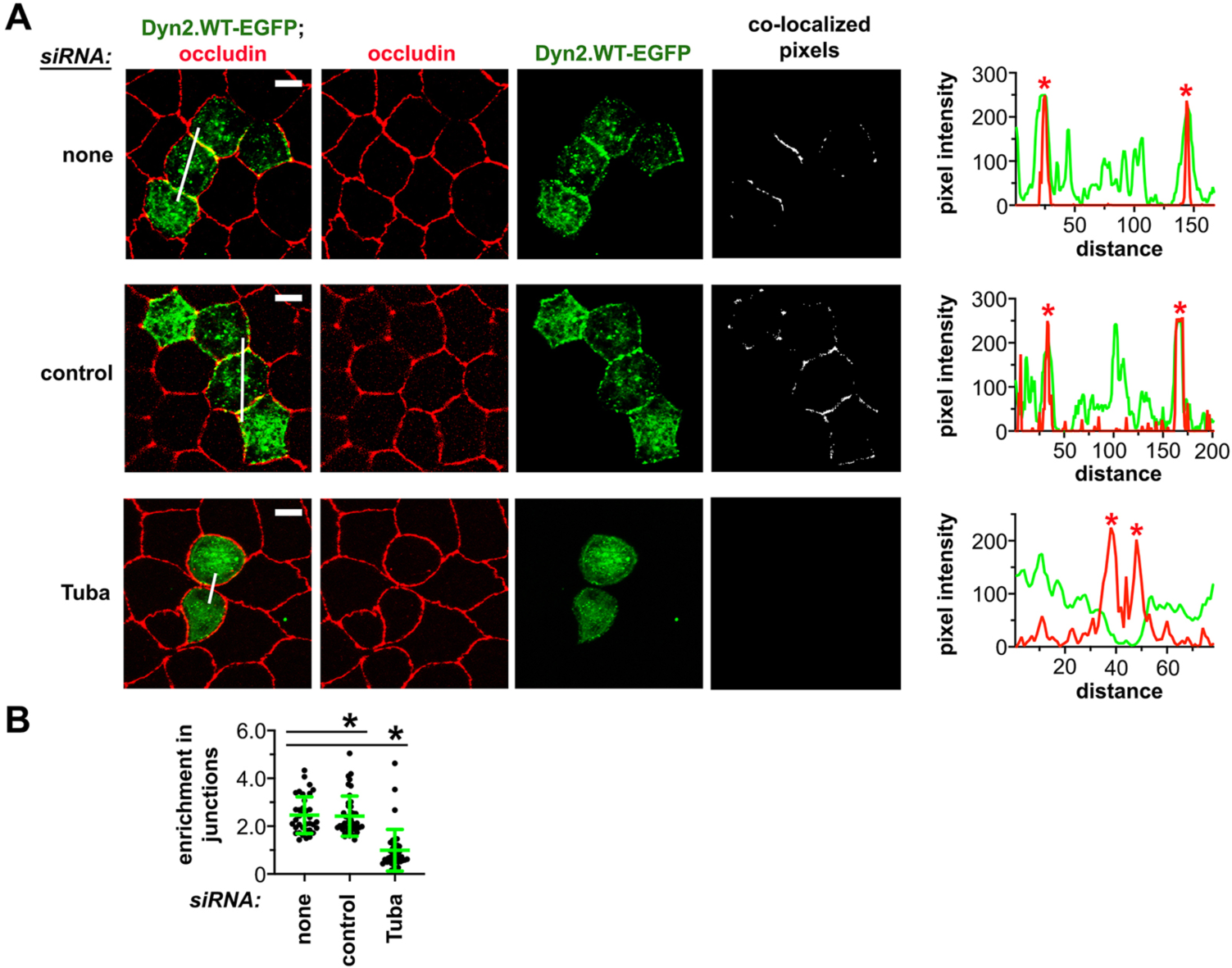
Localization of Dynamin 2 to the AJC depends on Tuba. A. Confocal microscopy images of Dynamin 2.WT-EGFP and occludin labeling in Caco-2 BBE1 cells subjected to control conditions or transfected with an siRNA targeting Tuba. The white lines in the left image panels show the regions used to generate the plot profile graphs on the right. Asterisks indicate cell junctions. Scale bars are 10 micrometers. B. The graph contains values for enrichment of Dynamin 2.WT-EGFP at cell junctions. Approximately 40 junctions were analyzed for each condition. *, P < 0.05.

In order to determine if Dynamin 2 contributes to cortical tension at the AJC, we used siRNAs to deplete Dynamin 2 or Tuba, and examined the effects on tight junction morphology in Caco-2 BBE1 cells. Under control conditions involving mock transfection or transfection with control siRNA, most tight junctions had a linear morphology, consistent with these structures experiencing cortical tension (Fig. 9A part i) (12, 15). As previously observed, transfection with an siRNA targeting Tuba caused a substantial proportion of cells to exhibit curved junctions, suggesting diminished cortical tension (Fig. 9 A part i) (12, 15). A similar result was caused by transfection with two different siRNAs that deplete Dynamin 2. The effects of siRNAs targeting Tuba or Dynamin 2 on junctional morphology were quantified as linear index values, which indicate the extent to which junctions differ from perfect linearity (12, 15, 51). Compared to control conditions, cells transfected with siRNAs against Dynamin 2 or Tuba had higher mean linear index values (Fig. 9A part ii). Similar results were observed for cells treated with dynasore (Fig. 9B). The effect of dynasore treatment resembled that produced by the compound Y27632, a Rho kinase inhibitor that relieves cortical tension by impairing the activity of non-muscle myosin II proteins (8, 11, 12, 52). The data in Figure 9A,B therefore provide evidence that Dynamin 2 and Tuba both contribute to cortical tension at the AJC.

**Figure 9.**
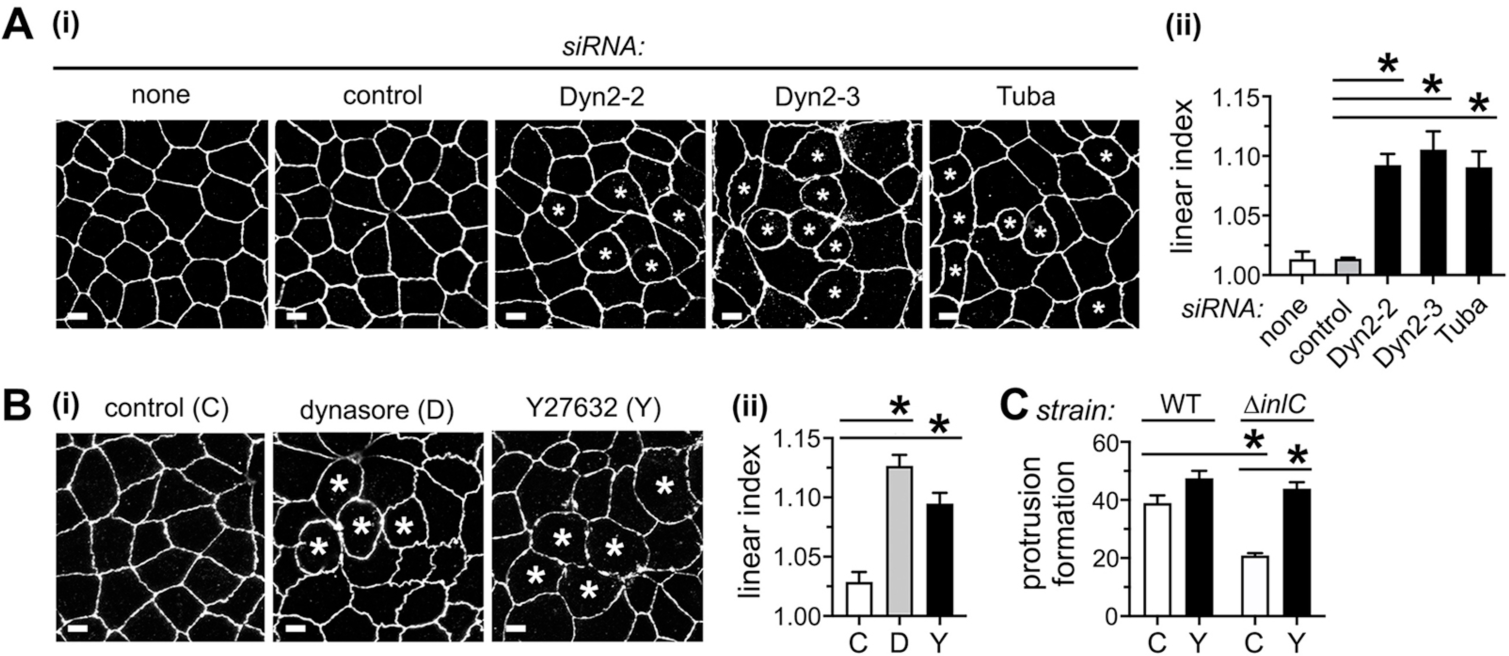
Genetic or pharmacological inhibition of Dynamin 2 or Tuba alters the structure of apical junctions. A. Effects of Dynamin 2 or Tuba RNAi on tight junction morphology. (i). Shown are representative confocal microscopy images of tight junctions in Caco-2 BBE1 cells subjected to control conditions or transfected with siRNAs against Dynamin 2 or Tuba. Tight junctions were detected using anti-occludin antibodies. Asterisks indicate cells with curved junctions. (ii). The graph contains linear index measurements from three experiments. In each experiment, between 95-127 junctions were analyzed per condition. B. Effect of treatment with dynasore or Y27632 on tight junctions. Images (i) and linear index values (ii) from three experiments are presented. In each experiment, between 62-68 junctions were assessed per condition. Scale bars indicate 10 micrometers. *, P <0.05, compared to the no siRNA or control siRNA conditions. C. Ability of Y27632 treatment to restore normal protrusion formation of the Δ*inlC* strain of *L. monocytogenes*. The data are mean +/- SEM values from three experiments. *, P < 0.05.

Previous results indicate that infection of Caco-2 BBE1 cells with WT *Listeria* causes tight junctions to slacken (15, 16, 43), similarly to the effects of depletion of Dynamin 2 or Tuba shown in Figure 9A,B. By contrast, infection with the Δ*inlC* mutant of *Listeria* fails to affect tight junction structure, indicating a key role for InlC in perturbing the AJC (15, 16, 43). We found that treatment of Caco-2 BBE1 cells with Y27632, which alters AJC morphology by inhibiting myosin II-mediated cortical tension (8, 11, 52) (Fig. 9B), restored normal protrusion formation to the Δ*inlC* mutant of *Listeria* without affecting protrusion efficiency of the WT strain (Fig. 9C). These effects of Y27632 were similar to those caused by depletion of Dynamin 2 or Tuba (Figs. 3A and 4B), suggesting that these two host proteins likely control *Listeria* spread through effects on cortical tension.

## DISCUSSION

Polarized epithelia, including Caco-2 BBE1 cells used in our studies, develop AJCs that are subjected to actomyosin-mediated cortical tension (8, 11, 12, 15, 53). Collectively, our results indicate that Dynamin 2 and Tuba act together to produce cortical tension that controls cell-to-cell spread by *Listeria*. Biochemical experiments showed that the SH31-4 region in Tuba binds directly to Dynamin 2 in a manner partly dependent on its PRD. Tuba and the PRD were necessary for recruitment of Dynamin 2 to tight junctions. Experiments involving RNAi or dynasore indicated that both Dynamin 2 and Tuba contribute to cortical tension and antagonize protrusion formation and spread of a *Listeria* strain deleted for the virulence gene *inlC* (Δ*inlC*).

Apart from Dynamin 2, at least two other human ligands of Tuba control spread of *Listeria* — the actin regulatory protein N-WASP and the GTPase Cdc42 (15, 16, 43) (Fig. 10). The SH36 domain in Tuba binds directly to N-WASP (16, 17). This SH36/N-WASP interaction promotes cortical tension and restrains protrusion formation by the Δ*inlC* mutant strain of *Listeria* (12, 15, 16). The DH domain in Tuba activates Cdc42 (12, 17), which also contributes to cortical tension and limits spread of the Δ*inlC* strain (43). By contrast, WT *Listeria* antagonizes Tuba/N-WASP complexes and inhibits Cdc42 activity, thereby alleviating tension and augmenting bacterial protrusion formation (15, 43). Antagonism of Tuba/N-WASP complexes is achieved by InlC, which binds directly to the SH36 domain in Tuba and displaces N-WASP from this domain (15, 16). The *Listeria* factor that downregulates Cdc42 remains unidentified (43). An interesting question is whether *Listeria* has a mechanism to antagonize Dynamin 2, or whether blocking interaction with N-WASP and inhibiting Cdc42 activity are the sole ways that this bacterium impairs Tuba function. InlC seems unlikely to affect interaction of Dynamin 2 with Tuba, since this bacterial protein does not bind to purified SH31-4 from Tuba (15) or to purified Dynamin 2 (our unpublished results). Therefore, if *Listeria* factors targeting Dynamin 2 exist, they are currently unknown.

**Figure 10.**
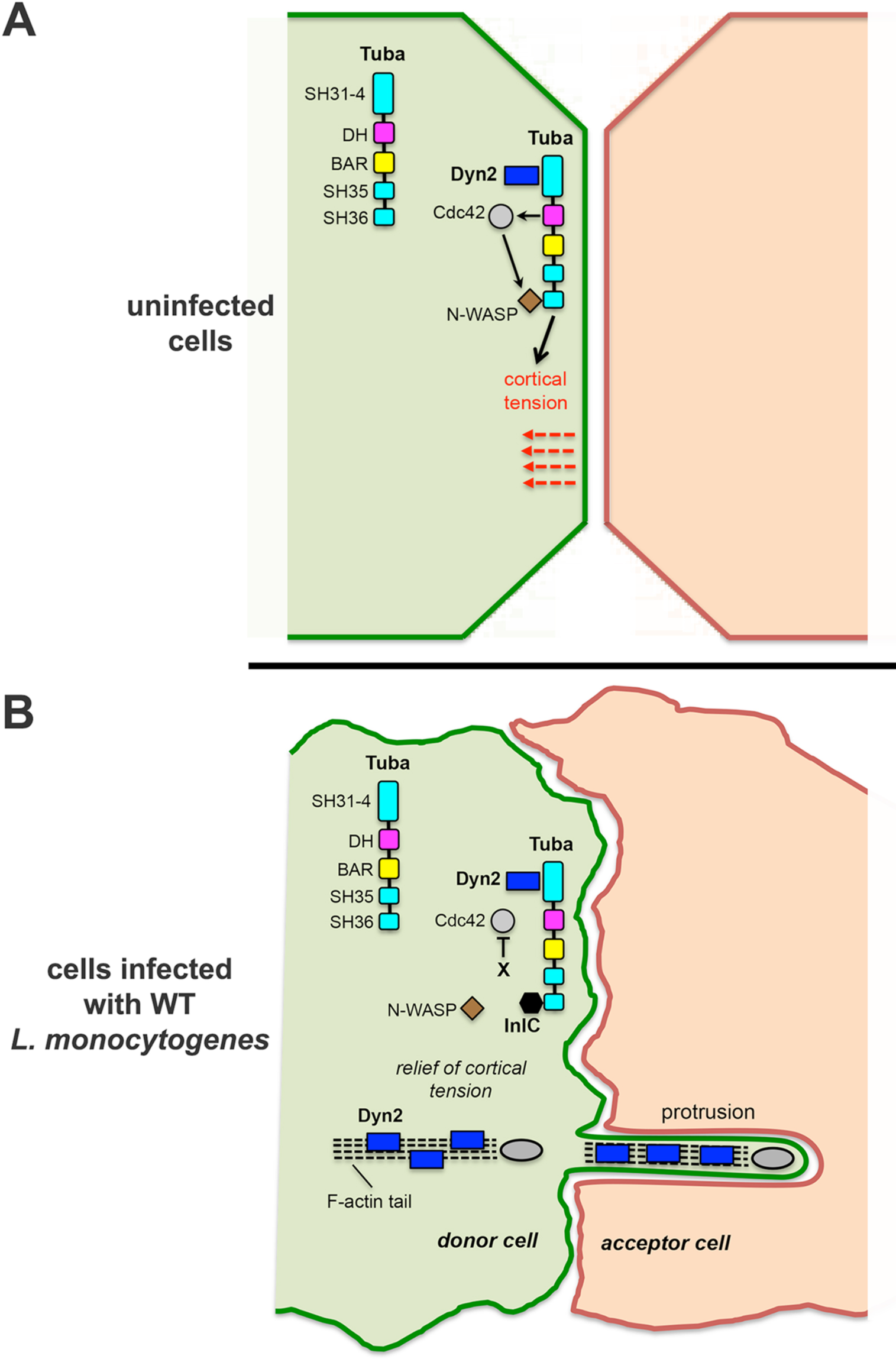
Model for role of Dynamin 2 in intercellular spread of *L. monocytogenes*. A. In uninfected Caco-2 BBE1 cells, Tuba recruits Dynamin 2 to the plasma membrane at the AJC. There, Tuba and Dynamin 2 act together to promote cortical tension. Also involved in generating tension are N-WASP, which binds to the SH36 domain in Tuba, and Cdc42 which is activated by the DH domain in Tuba (12, 17). Activated Cdc42 relieves autoinhibition of N-WASP, resulting in increased actin polymerization involved in cortical tension. B. In Caco-2 BBE1 cells infected with WT *L. monocytogenes*, at least two mechanisms interfere with the ability of Tuba and its ligands to promote cortical tension. First, InlC binds to SH36, thereby displacing N-WASP from this domain (15, 16). Secondly, an unidentified bacterial factor (‘X’) attenuates Cdc42 activity (43). Whether *L. monocytogenes* produces factors that antagonize Dynamin 2 is presently unknown. Also depicted in this diagram is the localization of Dynamin 2 to F-actin tails and in protrusions in host cells that donate these structures.

Interestingly, our results indicate that Dynamin 2 is recruited to *Listeria* protrusions in Caco2 BBE1 cells donating these structures, but not in cells accepting protrusions (Fig. 10). This recruitment to protrusions depended on Tuba and the Dynamin 2 PRD and was enhanced by the bacterial protein InlC. Since the Tuba and the PRD were needed to mobilize Dynamin 2 to junctions as well as to protrusions, we considered the possibility that accumulation of Dynamin 2 in protrusions might be a passive consequence of bacteria targeting tight junctions during cell-to-cell spread. *Listeria* and *Shigella flexneri*, another bacterium that undergoes cell-to-cell spread, are known to generate protrusions at tight junctions in some epithelial cell lines (54). However, in our studies with Caco-2 BBE1 cells, the tight junction protein occludin was not detected in protrusions made by WT or Δ*inlC* strains of *Listeria* (our unpublished data). The origin of Dynamin 2 in protrusions is therefore uncertain. It is also unclear what role protrusion-associated Dynamin 2 might play in spread of *Listeria*, given our finding that Tuba and Dynamin 2 control spread by modulating cortical tension.

In addition to accumulating in protrusions, Dynamin 2 was mobilized to *Listeria*-induced actin comet tails in the cell body in Caco-2 BBE1 cells (Fig. 10). Similar to the situation with protrusions, Dynamin 2 recruitment to actin comet tails required the PRD. However, neither Tuba nor InlC contributed to accumulation of Dynamin 2 in comet tails, indicating differences in the mechanisms of recruitment to protrusions and actin tails in the cell body. The PRD mediates direct interaction of Dynamin 2 with actin filaments (27), and indirect interaction through the actin regulatory protein cortactin (55, 56). These direct and/or indirect mechanisms may be responsible for mobilization of Dynamin 2 to comet tails during actin-based motility. Our results show that depletion of Dynamin 2 or treatment of cells with dynasore does not affect the proportion of bacteria that recruit F-actin in the cell body or the lengths of actin comet tails. These findings indicate that Dynamin 2, although present in comet tails, does not play a critical role in actin-based motility in the cytosol. Instead, the predominant role for Dynamin 2 in controlling spread of *Listeria* is at the level of protrusion formation.

Like *Listeria*, the bacterial pathogens *Shigella flexneri* and members of the spotted fever group (SFG) of *Rickettsia* undergo actin-based motility and protrusion formation to spread in human tissues (14, 57). *Shigella* infects intestinal epithelial cells, to produce inflammation and bloody diarrhea (58), whereas *Rickettsia* SFG spp. invade endothelial cells to cause vascular dysfunction (59). Similarly to *Listeria*, both *Shigella* and *Rickettsia* have evolved strategies to disrupt cortical tension at the AJC. However, the mechanisms that these three bacteria employ to modulate tension differ (14, 57). *Shigella* uses the type III secretion translocon protein IpaC to target human β-catenin, a critical component of adherens junctions (60). This interaction attenuates cortical tension and enhances protrusion formation and spread by *Shigella*. *Rickettsia* secretes an effector protein called Sca4 that disrupts a protein complex containing human vinculin and the adherens junction protein α-catenin, resulting in decreased intercellular tension and enhanced internalization of protrusions (61). Given that Dynamin 2 localizes to the AJC where it controls tight junction structure (29–31), future studies might reveal that this GTPase controls spread of *Shigella* and/or *Rickettsia* similarly to *Listeria*.

## MATERIALS AND METHODS

### Bacterial strains, mammalian cell lines and media

The WT *Listeria monocytogenes* strain EGD and the isogenic Δ*inlC* strain with an in-frame deletion in *inlC* have been previously described (15, 62). These strains were grown in brain heart infusion (BHI; Difco) broth and prepared for infection as detailed (Ireton et al., 1999). The human enterocyte cell line Caco-2 BBE1 (ATCC, CRL-2102) and the human epithelial cell line HeLa (ATTC, CCL-2) were cultured in Dulbecco’s modified Eagle’s medium (DMEM) with 4.5 g of glucose per liter and 2 mM glutamine (catalog no. 11995-065; Life Technologies), supplemented with 10% fetal bovine serum (FBS). In the case of Caco-2 BBE1 cells, 10 µg per mL of human transferrin (Sigma-Aldrich) was included in the growth medium. HeLa cells were grown in standard tissue culture plasticware, whereas Caco-2 BBE1 cells were cultured on transwell permeable supports (Corning 3450; 0.4 µm pore size), unless otherwise stated. Cell growth and bacterial infections were performed at 37°C under 5% CO2.

### Antibodies, inhibitors, and purified proteins

Rabbit antibodies used were anti-Dynamin 2 (Ab65556; Abcam), anti-glutathione-S-transferase (GST) (G7781; Sigma-Aldrich), anti-InlC (15), anti-*Listeria* (BD223021; Difco) (9272; Cell Signaling Technology), anti-Tuba SH31-4 (15), and anti-Tuba SH36 (15). Mouse monoclonal antibodies used were anti-GFP clones 7.7 and 13.1 (11814460001; Sigma-Aldrich), anti-hemagglutinin (HA) (MMS-101P; Covance), anti-occludin (331500; Invitrogen), and anti-tubulin (T5168; Sigma-Aldrich). Horseradish peroxidase (HRPO)-conjugated secondary antibodies were purchased from Jackson Immunoresearch. Secondary antibodies or phalloidin coupled to Alexa Fluor 488, Alexa Fluor 555, or Alexa Fluor 647 were obtained from Life Technologies. The inhibitors dynasore or Y27632 were procured from Sigma-Aldrich.

### Plasmids

Mammalian expression vectors used were pcDNA3.1-Ha-Dynamin 2.WT (gift of S. Schmid; Addgene 34684), pcDNA3.1-Ha-Dynamin 2.K44A (gift of S. Schmid, Addgene 34685), pEGFPC1 (Clontech), pEGFP-actin (15), pEGFP-Dynamin 2.K44A (gift of P. De Camilli; Addgene 22301), and pEGFP-Dynamin 2ΔPRD (50) (gift of P. De Camilli). The plasmid pEGFP-Dynamin 2.WT was constructed using a Quik Change II XL (Agilent) site-directed mutagenesis kit to change the GCC (alanine) codon at position 44 to AAG (lysine). The primers 5’-CAGAGCGCCGGCAAGAGTTCGGTGCTC-3’ and 5’ GAGCACCGAACTCTTGCCGGCGCTCTG-3’ were used for mutagenesis. Plasmid constructs allowing expression of glutathione S-transferase (GST)-tagged proteins containing InlC or the SH31-4, SH35, or SH36 regions in human Tuba were previously described (15, 17). Plasmids expressing GST-tagged WT Dynamin 2 or Dynamin 2 ΔPRD were constructed using the Takara Biosciences In-Fusion system and a plasmid encoding rat WT Dynamin 2 (Addgene 34684) as a template. The primers dyn2wtF (5’- TGGATCCCCGGAATTCATGGGCAACCGCGGGATGG-3’) and dyn2wtR (5’- GGCCGCTCGAGTCGACCTAGTCGAGCAGGGACGGC-3’) were used to PCR amplify Dynamin 2.WT, whereas dyn2wt F and dyn2dPRD (5’- GGCCGCTCGAGTCGACCTACGTGGACACGGTGCTGG-3’) were used for amplification of a fragment encoding Dynamin 2ΔPRD. The PCR products were subcloned into EcoRI/SalI sites of pGEX-4-T-1 and the correct DNA sequence was verified by sequencing.

### Purification of proteins

GST fusion proteins were expressed in *E. coli* strain BL21 λDE3 and purified as previously described (63, 64). Where appropriate, GST tags were removed by incubating 0.70 mg of protein coupled to glutathione Sepharose 4B (Cytiva) with 20 NIH units of thrombin (Sigma-Aldrich; 10602400001) overnight at room temperature. Protein lacking the GST tag was recovered in the supernatant following centrifugation at 6000 rpm for 5 min.

### siRNAs

The sequences of short interfering RNAs (siRNAs) used were 5’- CCCUCAAGGAGGCGCUCAAtt-3’ (Dyn2-2), 5’-CCAACAUGGACCUGGCCAAtt-3’ (Dyn2-3), and Tuba (5’-CAGAATCATGATGAGGCCAtt-3’. These siRNAs were obtained from Sigma-Aldrich (Dyn2-2 and Dyn2-3) or Qiagen (Tuba; catalog no. SI04220237). The negative, non-targeting control siRNA molecule #1 (catalog no. D-001210-01) was purchased from Dharmacon. This siRNA has two or more mismatches with all sequences in the human genome, indicating that it should not target host mRNAs.

### Transfection with siRNAs or plasmid DNA

For reverse transfection of Caco-2 BBE1 cells with siRNAs, 2.8 x10^5^ cells were seeded in transwells in the presence of 100 nM siRNA. Transfections were performed using Lipofectamine RNAiMax (13778075; Thermo Fisher Scientific), according to the manufacturers’ instructions. Plasmid DNA transfection was done using Lipofectamine 2000 (11668030; Thermo Fisher Scientific) as previously described (43). HeLa cells grown in 6-well plates were transfected with plasmid DNA expressing EGFP-tagged Dynamin 2.WT or Dynamin 2 ΔPRD using Lipofectamine 2000 as previously detailed (65).

### Co-precipitation experiments

For experiments using lysates of Caco-2 BBE1 cells (Fig. 1B,C,D and Fig. 2D), approximately 2 x 10^6^ cells were seeded in 10 cm plates and grown for four days prior to solubilization in lysis buffer (25 mM HEPES, pH 7.5, 150 mM NaCl, 1.0% IGEPAL CA-630, 1 mM EDTA, 1 mM sodium orthovanadate, 10 mM NaF, 1 mM phenylmethylsulfonyl fluoride [PMSF], and 10 µg per mL each of aprotinin and leupeptin) for 45 min at 4°C. Insoluble material was removed by centrifugation at 12,000 rpm at 4°C for 10 min. A preclearance step was performed by incubating ∼ 1.0 mg of lysate with 30 µg of GST alone on a rotating wheel at 4°C for 45 min. After centrifugation at 6000 rpm for 5 min, lysates were transferred to tubes containing 30 µg of GST-tagged SH31-4, SH35, SH36, Dynamin 2.WT, Dynamin 2ΔPRD, or GST alone. Samples were incubated with rotation at 4°C for 2 h followed by two washes in lysis buffer and two washes in lysis buffer with 0.20% IGEPAL-630. For studies with EGFP-tagged Dynamin 2.WT or Dynamin 2ΔPRD expressed in HeLa cells (Fig. 2C) ∼ 2 µg of GST fusion proteins were added to lysates. In experiments to assess binding of purified proteins (Fig. 2E), 3 nM of SH31-4 and 30 nM of GST-tagged Dynamin 2.WT or Dynamin 2ΔPRD proteins were incubated in binding buffer (20 mM Hepes pH 7.4, 150 mM NaCl, 0.1% Tween 20, 0.5% bovine serum albumin [BSA], 2 mM each of CaCl_2_ and MgCl_2_, 1 mM PMSF, and 10 µg of aprotinin and leupeptin) for 2 hours at 4°C. Samples were washed twice in binding buffer and twice in wash buffer, which lacked Tween 20. For all co-precipitation experiments, pellets obtained from the last centrifugation were boiled in sample buffer prior to migration on SDS-polyacrylamide gels for Western blotting.

### Western blotting

Western blotting studies involved migration on 7.5%, 9.0%, 10.0%, 12.0%, or 15.0% SDS-polyacrylamide gels, transfer of samples to PVDF membranes, incubation with primary antibodies or secondary antibodies coupled to horse radish peroxidase, and detection using Clarity Max Western ECL substrate (Bio-Rad), as described previously (66). Chemiluminescence was imaged using an Odyssey imaging system (Li-Cor Biosciences). Bands in Western blot images were quantified using ImageJ software as described (42). For experiments to confirm target protein depletion after siRNA transfection (Figs. 3Ai and 4A), Caco-2 BBE1 cells were solubilized in radioimmunoprecipitation assay (RIPA) buffer (1% Triton X-100, 0.25% sodium deoxycholate, 0.05% SDS, 50 mM Tris-HCl [pH 7.5], 2 mM EDTA, 150 mM NaCl, 1 mM PMSF, and 10 µg per mL each of aprotinin and leupeptin) approximately 72 h post-transfection. Protein concentrations of lysates were determined using a bicinchoninic acid (BCA) assay kit (Pierce), and equal protein amounts of each sample were migrated on gels prior to transfer to PVDF membranes, antibody incubation, and detection with ECL. For co-precipitation experiments (Figures 1 and 2), the GST proteins in precipitates were detected by stripping membranes in buffer containing 7.5 g/L glycine, 0.10% Triton X-100, 0.10% SDS (pH 2.3) and stained using Coomassie Brilliant Blue R-250 solution.

### Confocal microscopy imaging

An inverted Olympus FV1200 laser scanning confocal microscope equipped with 40x dry and 60x 1.35 NA oil immersion objectives, laser lines of 405, 488, 543, and 633 nm, and photomultiplier tube detectors was used for imaging studies. This work included analysis of bacterial protrusion formation and cell-to-cell spread (Figs. 3, 4, S2, S3, S4), recruitment of Dynamin 2 to protrusions or F-actin tails (Figs. 5, 6, S5, S6, S7), endocytosis of transferrin (Fig. 3D), localization of proteins to apical junctions (Figs. 7 and 8), and morphology of tight junctions (Fig. 9).

### Protrusion formation assays

For experiments involving siRNA-mediated depletion of Dynamin 2 or Tuba (Figs. 3A and 4A,B), Caco-2 BBE1 cells in transwells were reverse transfected with siRNAs for ∼ 48 h and then transfected with plasmid DNA expressing EGFP-actin. About 24 h later, cell monolayers were infected with a ∼ 10:1 ratio of WT or Δ*inlC* strains of *L. monocytogenes* for 1.5 h in DMEM lacking gentamicin. Cells were washed in DMEM and incubated for another 4.5 h in the same medium containing 20 µg per mL gentamicin. Cells were then washed twice in PBS, fixed in PBS containing 3% paraformaldehyde, quenched in PBS with 50 mM ammonium chloride, and permeabilized in PBS with 0.4% Triton X-100. Samples were labeled with rabbit anti-*L. monocytogenes* antibodies, anti-rabbit-Alexa Fluor 647 secondary antibodies, and phalloidin Alexa Fluor 555 to detect F-actin. Slides were mounted using Molwiol supplemented with DABCO (1,4-diazabicylo[2.2.2]octane), as previously described (15, 41). Experiments involving treatment with dynasore or Y27632 (Figs. 3D and 9C) were performed similarly to the siRNA studies, except that Caco-2 BBE1 cells were infected for 1.5 h in the absence of drug, followed by incubation for another 4.5 h in DMEM containing 80 µM dynasore or 25 µM Y27632 and gentamicin. DMEM containing 0.20% dimethyl sulfoxide (DMSO) was used as a control for the dynasore experiments. Since the Y27632 stock solution (50 mM) was dissolved in sterile PBS, DMEM alone served as the control condition.

Protrusion efficiency was measured using an established approach that takes advantage of the observation that approximately 25% of the transfected population of cells expresses EGFP-actin. This allows protrusions to be detected as structures projecting from host cells expressing EGFP-actin into surrounding cells lacking EGFP-actin (15, 16, 41, 43). Samples were imaged by confocal microscopy using a 63x objective. Images from serial sections spaced 1.0 µm apart were acquired and used to quantify three different types of actin structures associated with bacteria. These structures were protrusions, F-actin tails in the cell body, and bacteria labeled with symmetric actin (Fig. S2). Protrusions were identified as *L. monocytogenes* with F-actin tails emanating from EGFP-positive cells into EGFP-negative cells. Actin tails in cell bodies were defined as being located within the boundaries of EGFP-actin-positive cells, and symmetric actin was classified as structures coating the entire surface of bacteria. For each experiment, approximately 50 protrusions were scored for the WT *L. monocytogenes* strain in cells subjected to control conditions. These controls were mock transfection in the absence of siRNA (Figs. 3A or 4B), incubation in DMEM with 0.20% DMSO (Fig. 3D) or incubation in DMEM alone (Fig. 9C). Corresponding numbers of total actin structures associated with bacteria (i.e. protrusions, actin tails, and symmetric actin) were quantified for all other conditions. Protrusion efficiency was expressed as the percentage of total bacterial actin-associated structures present in protrusions. For studies with Ha-tagged WT or K44A forms of Dynamin 2 (Fig. 3C), protrusion formation was quantified similarly to experiments with EGFP-actin, except that the signal from Ha labeling was used to identify protrusions projecting from transiently transfected cells.

### Quantification of F-actin recruitment and actin tail lengths

Analyses of F-actin recruitment by bacteria and lengths of actin tails in cells treated with siRNAs against Dynamin 2 or Tuba, expressing Ha-tagged WT and K44A forms of Dynamin 2, or treated with dynasore or Y27632 (Fig. S4) were performed on the same samples used for quantification of protrusion formation (Figs. 3A, 4B, 4D, and 9C). F-actin recruitment efficiencies were defined as the percentage of total bacteria in the main body of the host cell that had F-actin tails or symmetric actin. The lengths of F-actin tails were measured using Image J (version 1.54f) software, as described (15, 16, 41).

### Cell-to-cell spread

Caco-2 BBE1 cells in transwells were mock transfected in the absence of siRNA, transfected with a control siRNA, or transfected with siRNAs targeting Dynamin 2 or Tuba. Approximately 72 h post-transfection, cells were infected with WT or Δ*inlC* strains of *L. monocytogenes* using a multiplicity of infection (MOI) of ∼ 10:1. Cells were infected for 1.5 h in DMEM lacking gentamicin, washed, and then incubated for another 10.5 h in DMEM with 20 µg/mL gentamicin. Cells were then washed and fixed in PBS containing 3% PFA. Samples were quenched by incubation in PBS with 50 mM ammonium chloride, permeabilized using PBS containing 0.40% Triton X-100, and labeled with rabbit anti-*Listeria* antibodies, anti-rabbit Alexa Fluor 647 secondary antibodies, and phalloidin coupled to Alexa Fluor 555 to detect F-actin.

Spreading efficiencies of *L. monocytogenes* strains were quantified by measuring the sizes of regions of infected cells, as described (16, 41, 43). Confocal microscopy images were acquired using a 40x objective. ImageJ software was used to measure surface areas of regions of contiguous infected human cells, referred to as ‘foci’ (Fig. S3). Approximately 25 foci were measured for each condition in each of three experiments.

### Analysis of recruitment of Dynamin during infection

For experiments involving EGFP-tagged Dynamin 2 proteins (Figs. 5, 6, S7, and S8), Caco-2 BBE1 cells were transfected with plasmids expressing EGFP-tagged WT Dynamin 2 or Dynamin 2ΔPRD. About 24 h after transfection, cells were infected with WT or Δ*inlC* strains of *L. monocytogenes* at an MOI of ∼ 10:1 for 1.5 h in the absence of gentamicin followed by incubation for another 4.5 h in the presence of gentamicin. Samples were then fixed, permeabilized, labeled, and mounted on slides as described for the protrusion formation assays. In experiments involving siRNA-mediated depletion of Tuba (Figs. 6 and S8), Caco-2 BBE1 cells were reversed transfected with siRNAs approximately 48 h prior to plasmid DNA transfection.

Samples were imaged by confocal microscopy similarly to as described for the protrusion formation studies. Protrusions were identified as F-actin tails projecting from cells expressing EGFP-tagged WT Dynamin 2 or Dynamin 2ΔPRD into neighboring cells lacking EGFP protein expression. The threshold function of ImageJ software was used to determine fold enrichment (FE) values for each protrusion similarly to as previously described (41). FE is defined as the mean pixel intensity of tagged Dynamin 2 protein in protrusions normalized to the mean pixel intensity throughout the entire infected human cell. The numbers of FE values determined for each condition are provided in the legends to Figures 5 and 6.

Experiments assessing the recruitment of endogenous Dynamin 2 to *L. monocytogenes* protrusions (Fig. S5) were performed similarly to those described above, with the following differences. First, Caco-2 BBE1 cells in transwells were transfected with a plasmid expressing EGFP in order to detect protrusions. Secondly, infected samples were fixed in PBS with 3% PFA, permeabilized with 0.40% Triton X-100, and then labeled by incubation with a rabbit anti-Dynamin 2 antibody and anti-rabbit antibodies conjugated to Alexa Fluor 555. F-actin was labeled using phalloidin coupled to Alexa Fluor 647. FE values for ∼ 50 protrusions of WT or Δ*inlC* strains were quantified.

For studies on the localization of EGFP-tagged Dynamin 2 in host cells donating or accepting protrusions (Fig. S6), the plot profile function of ImageJ software was used to obtain line scans of pixel intensities of Dynamin 2.WT-EGFP and F-actin. Peaks of F-actin were used to identify protrusions. Average pixel intensities of Dynamin 2.WT-EGFP in regions within and outside of the F-actin peaks were determined and the fold increase in Dynamin 2 was quantified as the ratio of these values.

### Quantification of localization of EGFP-tagged WT or ΔPRD Dynamin 2 to F-actin tails or symmetric actin

The studies in Figures S7 and S8 were performed similarly to protrusion recruitment experiments described above, except that FE values for F-actin comet tails or symmetric actin within the main body of infected human cells were quantified. The numbers of FE determinations are given in the figure legends.

### Transferrin endocytosis assay

Caco-2 BBE1 cells in transwells were incubated in DMEM lacking FBS and transferrin for 2 h, followed by treatment with 80 µM dynasore or 0.20% of the vehicle DMSO for 4.5 h. Human transferrin conjugated to Alexa Fluor 647 (20 µg/mL) was then added to both the top and bottom chambers of the transwells, and cells were incubated at 37°C in 5% CO2 for 30 min to allow endocytosis. Cells were washed twice in ice-cold PBS with 1 mM CaCl_2_, 1 mM MgCl_2_, and 0.10% BSA. External transferrin bound to the surface of cells was removed by acid stripping for 2 min in ice-cold 100 mM MES, pH 5.5. Cells were washed three times in PBS with 1 mM CaCl_2_ and 1 mM MgCl_2_ and fixed in the same PBS solution with 3% PFA for 20 min. Samples were then quenched in PBS with 50 mM ammonium chloride, permeabilized in PBS with 0.40% Triton X-100, and labeled with phalloidin coupled to Alexa Fluor 555.

For quantification of transferrin endocytosis, serial sections of samples spaced 1 µm apart were imaged by confocal microscopy using a 63x objective. Laser power and detector settings were kept identical for imaging of all samples. ImageJ software was used to measure mean pixel intensities of transferrin-Alexa Fluor 647 in z-project images. Approximately 5 fields of view were imaged for each condition in each of three experiments.

### Localization of Dynamin 2 and Tuba in uninfected cells

For assessment of localization of EGFP-tagged WT Dynamin 2 or Dynamin 2ΔPRD at tight junctions (Fig. 7A), Caco-2 BBE1 cells were transfected with plasmids expressing EGFP-tagged proteins for ∼ 48 h, followed by fixation in cold methanol. Cells were labeled with mouse anti-occludin antibodies and anti-mouse antibodies conjugated to Alexa Fluor 555. Samples were imaged by confocal microscopy using a 63x objective. Co-localized pixel analysis of EGFP-tagged Dynamin 2 proteins and the tight junction protein occludin was performed using the co-localization threshold plug-in of ImageJ. The plot profile function of ImageJ was used to assess the spatial co-distribution of these proteins. Experiments analyzing localization of Tuba and EGFP-tagged Dynamin 2 proteins (Fig. 7C) were performed similarly to as described above, except that cells were labeled with rabbit antibodies recognizing the SH36 domain of Tuba (15). For studies of localization of endogenous Dynamin 2 or Tuba to tight junctions (Fig. 7B), rabbit antibodies against Dynamin 2 or the Tuba SH36 domain were used. Experiments assessing the effect of an siRNA targeting Tuba on localization of EGFP-tagged WT Dynamin 2 (Fig. 8) were performed similarly to those in Figure 7A, except that Caco-2 BBE1 cells were transfected with siRNAs approximately 48 h prior to transfection with plasmid DNA. The enrichment of Tuba in each tight junction imaged was quantified by measuring the mean pixel intensity of Tuba in the region that overlaps occludin labeling and the mean pixel intensity of Tuba outside of this region. Enrichment was quantified as the ratio of these two values.

### Assessment of tight junction morphology

Caco-2 BBE1 cells in transwells were fixed in cold methanol approximately 72 h after reverse transfection with siRNAs targeting Dynamin 2 or Tuba (Fig. 9A). Tight junctions were labeled using mouse anti-occludin antibodies and anti-mouse antibodies coupled to Alexa Fluor 488. Experiments with dynasore or Y27632 (Fig. 9B) were performed similarly, except that cells were treated with 80 µM dynasore or 50 µM Y27632 for 4.5 h prior to fixation. After confocal microscopy imaging, 60-70 cell junctions for each condition in each of three experiments were analyzed for linear index values. Linear indices were determined by using ImageJ software to measure the distance of a junctional trace and also of a straight line between the junctional vertices, as described (12, 15, 16, 42, 43, 51). The linear index of a junction was calculated as the ratio between these two values. Approximately 40 enrichment values were quantified for each condition.

### Statistical analysis

Statistical analysis was performed using Prism (version 9.3.1; GraphPad Software). In comparisons of data from three or more conditions, analysis of variance (ANOVA) was used. The Tukey-Kramer test was used as a posttest. For comparisons of two conditions, an unpaired t test was performed. A P-value of 0.05 or lower was considered significant.

## ACKNOWLEDGEMENTS

We thank Roman Mortuza for performing site-directed mutagenesis. Nicole Lee is a recipient of a University of Otago doctoral scholarship. This work was supported by grants from the University of Otago Research Committee, the Health Research Council of New Zealand (22/296), and the Marsden Fund of the Royal Society of New Zealand (22-UOO-098) awarded to K.I.

## FIGURE LEGENDS

**Figure S1. Purified proteins used for co-precipitation studies.** Proteins were migrated on 9%, 10%, or 12% SDS-polyacrylamide gels and stained using Coomassie Blue or Western blotted as indicated. A. GST fusion proteins used for precipitations in Figure 1B. B. GST-tagged proteins used for experiments in Figures 1C and 2C. C. GST-tagged Dynamin 2 or InlC proteins used for studies in Figures 1D, 2D, and 2E. (i) Coomassie Blue-stained gels with GST-tagged full-length Dynamin 2 wild-type (Dyn2.WT), Dynamin 2ΔPRD (Dyn2ΔPRD), or InlC. (ii). GST-Dynamin 2 proteins subjected to Western blotting with antibodies recognizing GST or the PRD in Dynamin 2. The arrows indicate full-length GST-Dyn2.WT or GST-Dyn2ΔPRD. The remaining bands are likely degradation products, based on their reactivity to anti-GST antibodies. Similar degradation products of purified GST-Dyn2.WT were previously observed by others (68). D. SH31-4 and SH36 proteins used for binding studies in Figure 2E,F. Also included are GST-SH31-4 and GST-SH36 prior to removal of the GST tag. The amounts of each protein loaded are 1.0 µg each of GST-SH31-4 and SH31-4, 1.0 µg of GST-SH36, and 1.0 or 2.0 µg of SH36.

**Figure S2. Bacterial-associated actin structures used for quantification of protrusion formation.** Shown are representative confocal microscopy images of actin structures used to quantify protrusion formation efficiency, as described in the Materials and Methods. Caco-2 BBE1 cells transiently expressing EGFP-actin are colored green, total F-actin in all cells is red, and bacteria are blue. Protrusions containing *L. monocytogenes* are identified as structures projecting from cells expressing EGFP-actin into surrounding cells lacking EGFP-actin. Bacteria undergoing actin-based motility have F-actin tails in the main body of host cells. Bacteria symmetrically decorated with F-actin represent a step preceding tail formation.

**Figure S3. Representative images of infection foci used to assess cell-to-cell spread.** Representative confocal microscopy images of foci of infection produced by wild-type (WT) or Δ*inlC* strains of *L. monocytogenes* are shown. These images are some of those used to determine mean focus size data in Figure 3B part i. The yellow lines delimit the borders of foci in a monolayer of infected Caco-2 BBE1 cells. The numerals in the right panels are the sizes of the foci in square micrometers. Scale bars indicate 20 micrometers.

**Figure S4. Lack of effect of inhibition of Dynamin 2 or Tuba on F-actin recruitment or actin-based motility of *L. monocyogenes*.** The graphs contain data on bacterial recruitment of F-actin or actin tail lengths in Caco-2 BBE1 cells treated with siRNAs against Dynamin 2 (A and B) or Tuba (C), transiently expressing Ha-tagged wild-type Dynamin 2 or Dynamin 2.K44A proteins (D), incubated with dynasore (E), or treated with Y27632 (F). These data are mean values +/- SEM from three experiments and were obtained from the same images used to quantify protrusion formation efficiencies in Figure 3A, 4B, and 9C. For each condition of each experiment, approximately 200-500 intracellular bacteria were evaluated for F-actin recruitment and 50-100 actin tail lengths were measured. No statistically significant differences were observed between any of the conditions. In part D, ‘NS’ has been added to emphasize lack of statistical significance, despite slightly lower mean values in cells expressing Ha-tagged K44A Dynamin 2.

**Figure S5. Recruitment of endogenous Dynamin 2 to *Listeria* protrusions.** A. Images of Caco-2 BBE1 cells infected with WT or Δ*inlC* strains of *L. monocytogenese* for 6h. The arrows indicate protrusions projecting from host cells expressing EGFP into neighboring cells lacking EGFP. Images of protrusions are expanded in panels on the right. Scale bars indicate 5 micrometers. B. The graph contains data of fold enrichment (FE values) for endogenous Dynamin 2 in protrusions. Approximately 50 FE values were obtained for both the WT and Δ*inlC* bacterial strains. *, P < 0.05.

**Figure S6. Dynamin 2 is not recruited in cells accepting protrusions.** A. Accumulation of Dynamin 2 around a protrusion in a host cell donating this structure. (i). An image of a protrusion made by WT *L. monocytogenes* projecting from a Caco-2 BBE1 cell expressing EGFP-Dynamin 2.WT into a host cell lacking this protein. The protrusion is indicated with an arrow. The scale bar is 10 micrometers. (ii). Plot profile measurements showing accumulation of Dynamin 2 around the protrusion. The white line indicates the region analyzed for fluorescence of F-actin and Dynamin 2. The graph displays the corresponding pixel intensity measurements of Dynamin 2 (green) or F-actin (red) over the line. The protrusion is the F-actin peak indicated with an asterisk. B. Lack of accumulation of Dynamin 2 around a protrusion in a recipient host cell. (i) Image of a protrusion projecting from a Caco-2 BBE1 cell lacking EGFP-tagged Dynamin 2.WT into an adjacent cell expressing this protein. The scale bar represents 10 micrometers (ii). A plot profile graph showing pixel intensities across the white line. C. Quantification of plot profile data. The graph displays the mean fold increase -/+ SEM in Dynamin 2.WT-EGFP around 25 donated or accepted protrusions. *, P < 0.05.

**Figure S7. Dynamin 2 accumulates around F-actin tails and bacteria in the host cell body.** A. Representative images of distribution of EGFP-tagged Dynamin 2.WT or Dynamin 2ΔPRD in Caco-2 BBE1 cells infected with WT or Δ*inlC* strains of *L. monocytogenes* for 6 h. Arrows indicate bacteria with F-actin tails or symmetrically distributed F-actin. Scale bars represent 10 micrometers. B. Quantification of fold enrichment (FE) values for tagged Dynamin 2 proteins in F-actin tails or symmetrically distributed F-actin. Data are means +/- SEM. For each condition, between 12-16 F-actin tails or 28-83 symmetric actin structures were analyzed. *, P < 0.05.

**Figure S8. Tuba does not affect localization of Dynamin 2 to structures associated with actin-based motility in the cytosol.** A. Representative confocal microscopy images of EGFP-tagged WT Dynamin 2 in infected Caco-2 BBE1 cells that were subjected to control conditions or transfected with an siRNA targeting Tuba. The left panels contain images of F-actin comet tails, whereas the right panels have images of bacteria decorated with symmetric actin. Dynamin 2.WT-EGFP is green, F-actin is red, and bacteria are blue. In the right panels, not all bacteria have blue labeling, perhaps due to F-actin interfering with antibody labeling. Scale bars are 10 micrometers. B. Quantified FE values for EGFP-Dynamin 2.WT in F-actin tails or around bacteria with symmetric F-actin. Data are means +/- SEM. For each condition, between 39-51 F-actin tails or 52-64 symmetric actin structures were analyzed. Statistically significant differences were not observed between any of the conditions.

## REFERENCES

1. Schlech WF. 2019. Epidemiology and clinical manifestations of *Listeria monocytogenes* infection. Microbiol Spectr 7.

2. Ireton K. 2013. Molecular mechanisms of cell-cell spread of intracellular bacterial pathogens. Open Biol 3:130079.

3. Kammoun H, Kim M, Hafner L, Gaillard J, Disson O, Lecuit M. 2022. Listeriosis, a model infection to study host-pathogen interactions in vivo. Curr Opin Microbiol 66:11–20.

4. Lamond NM, Freitag NE. 2018. Probing the balance between protection from pathogens and fetal tolerance. Pathogens 7:52.

5. Drolia R, Bhunia AK. 2019. Crossing the Intestinal Barrier via *Listeria* Adhesion Protein and Internalin A. Trends Microbol 27:408–425.

6. Rusu AD, Georgiou M. 2020. The multifarious regulation of the apical junctional complex. Open Biol 10:190278.

7. Vazquez-Boland JA, Kuhn M, Berche P, Chakraborty T, Dominguez-Bernal G, Goebel W, Gonzalez-Zorn B, Wehland J, Kreft J. 2001. Listeria pathogenesis and molecular virulence determinants. Clin Microbiol Rev 14:584–640.

8. Acharya BR, Wu SK, Lieu ZZ, Parton RG, Grill SW, Berdhadsky AD, Gomez GA, Yap AS. 2017. Mammalian diaphanous 1 mediates a pathway for E-cadherin to stablize epithelial barriers through junctional contractility Cell Rep 18:2854–2867.

9. Charras G, Yap AS. 2018. Tensile Forces and Mechanotransduction at Cell-Cell Junctions. Curr Biol 28:R445–R457.

10. Citi S. 2019. The mechanobiology of tight junctions. Biophys, Rev 11:783–793.

11. J.M. L, Gomez GA, Verma S, Moussa EJ, Wu SK, Priya R, Hoffman BD, Grashoff C, Schwartz MA, Yap AS. 2014. Tension-sensitive actin assembly supports contractility at the epithelial zonula adherens. Curr Biol 24:1689–1699.

12. Otani T, Ichii T, Aono S, Takeichi M. 2006. Cdc42 GEF Tuba regulates the junctional configuration of simple epithelial cells. J Cell Biol 175:135–146.

13. Cunningham KE, Turner JR. 2012. Myosin light chain kinase: pulling the strings of epithelial tight junction function. Ann NY Acad Sci 1258:34–42.

14. Dowd GC, Mortuza R, Ireton K. 2021. Molecular mechanisms of intercellular dissemination of bacterial pathogens Trends Microbol 29:127–141.

15. Rajabian T, Gavicherla B, Heisig M, Muller-Altrock S, Goebel W, Gray-Owen SD, Ireton K. 2009. The bacterial virulence factor InlC perturbs apical cell junctions and promotes cell-to-cell spread of *Listeria*. Nat Cell Biol 11:1212–1218.

16. Polle L, Rigano LA, Julian R, Ireton K, Schubert WD. 2014. Structural details of human Tuba recruitment by InlC of *Listeria monocytogenes* elucidate bacterial cell-cell spreading. Structure 22:304–314.

17. Salazar MA, Kwiatkowski AV, Pellegrini L, Cestra G, Butler MH, Rossman KL, Serna DM, Sondek J, Gertler FB, De Camilli P. 2003. Tuba, a novel protein containing Bin/Amphiphysin/Rvs and Dbl homology domains, links dynamin to regulation of the actin cytoskeleton. J Biol Chem 278:49031–49043.

18. Gonzalez-Jamett AM, Momboisse F, Haro-Acuna V, Bevilacqua JA, Caviedes P, Cardenas AM. 2013. Dynamin-2 function and dysfunction along the secretory pathway. Front Endocrinol 4:126.

19. Laiman J, Lin SS, Liu YW. 2023. Dynamins in human diseases: differential requirement of dynamin activity in distinct tissues. Curr Opin Cell Biol 81:102174.

20. Perrais D. 2022. Cellular and structural insight into dynamin function during endocytic vesicle formation: a tale of 50 years of investigation. Biosci Rep 42:BSR20211227.

21. Ferguson SM, Brasnjo G, Hayashi M, Wölfel M, Collesi C, Giovedi S, Raimondi A, Gong LW, Ariel P, Paradise S, O’toole E, Flavell R, Cremona O, Miesenböck G, Ryan TA, De Camilli P. 2007. A selective activity-dependent requirement for dynamin 1 in synaptic vesicle endocytosis. Science 316:570–574.

22. Raimondi A, Ferguson SM, Loux X, Armbruster M, Paradise S, Giovedi S, Messa M, Kono N, Takasaki J, Capello V, O’Toole E, Ryan TA, De Camilli P. 2011. Overlapping role of dynamin isoforms in synaptic vesicle formation. Neuron 70:1100–1104.

23. Cao H, Chen J, Awoniya M, Henley JR, McNiven MA. 2007. Dynamin 2 mediates fluid-phase micropinocytosis in epithelial cells. J Cell Sci 120:4167–4177.

24. Henley JR, Krueger WE, Oswald BJ, McNiven MA. 1998. Dynamin-mediated internalization of caveolae. J Cell Biol 141:85–99.

25. Lamaze C, Dujeancourt A, Baba T, Lo CG, Benmerah A, Dautry-Varsat A. 2001. Interleukin 2 receptors and detergent-resistant membrane domains define a clathrin-independent endocytic pathway. Mol Cell 7:661–671.

26. Menon M, Schafer DA. 2013. Dynamin: Expanding its scope to the cytoskeleton. Int Rev Cell Mol Biol 302:187–219.

27. Zhang R, Lee DM, Jimah JR, Gerassimov N, Yang C, Kim S, Luvsanjav D, Winkelman J, Mettlen M, Abrams ME, Kalia R, Keene P, Pandey P, Ravaux B, Kim JH, Ditlev JA, Zhang G, Rosen MK, Frost A, Alto NM, Gardel M, Schmid S, Svitkina TM, Hinshaw JE, Chen EH. 2020. Dynamin regulates the dynamics and mechanical strength of the actin cytoskeleton as a multifilament actin-bundling protein. Nat Cell Biol 22:674–688.

28. Yamada H, Takeda T, Michiue H, Abe T, Takei K. 2016. Actin bundling by dynamin 2 and cortactin is implicated in cell migration by stabilizing filopodia in human non-small lung carcinoma cells. Int J Oncol 49:877–886.

29. Chua J, Rikhy R, Lippincott-Schwartz J. 2009. Dynamin 2 orchestrates the global actomyosin cytoskeleton for epithelial maintenance and apical constriction. Proc Natl Acad Sci USA 49:20770–20775.

30. Eaton AF, Clayton DR, Ruiz WG, Griffiths SE, Rubio ME, Apodaca G. 2019. Expansion and contraction of the umbrella cell apical junctional ring in response to bladder filling and voiding. Mol Biol Cell 30:2037–2052.

31. Lynn KS, Easley KF, Martinez FJ, Reed RC, Schlingmann B, Koval M. 2021. Asymmetric distribution of dynamin-2 and beta-catenin relative to tight junction spikes in alveolar epithelial cells. Tissue Barriers 9:1929786.

32. Li J, Fujise K, Wint H, Senju Y, Suetsugu S, Yamada H, Takei K, Takeda T. 2021. Dynamin 2 and BAR domain protein PACSIN 2 cooperatively reslte formation and maturation of podosomes. Biochem Biophys Res Commun 571:145–151.

33. Ochoa GC, Slepnev VI, Neff L, Ringstad N, Takei K, Daniell L, Kim W, Cao H, McNiven MA, Baron R, De Camilli P. 2000. A functional link between dynamin and the actin cytoskeleton at podosomes. J Cell Biol 150:377–389.

34. Burton KM, Cao H, Cheng J, Qiang L, Krueger WE, Johnson KM, Bamlet WR, Zhang L, McNiven MA, Razidlo GL. 2020. Dynamin 2 interacts with alpha-catenin 4 to drive tumor cell invasion. Mol Biol Cell 31:439–451.

35. Zhang Y, Nolan M, Yamada H, Watanabe M, Nasu Y, Takei K, Takeda T. 2016. Dynamin 2 GTPase contributes to invadopodia formation in invasive bladder cancer cells. Biochem Biophys Res Commun 480:409–414.

36. Gray NW, Fourgeaud L, Huang B, Chen J, Cao H, Oswald BJ, Hemar A, McNiven MA. 2003. Dynamin 3 is a component of the postsynapse, where it interacts with mGluR5 and Homer. Curr Biol 13:510–515.

37. Kurklinsky S, Chen J, McNiven MA. 2011. Growth cone morphology and spreading are regulated by dynamin-cortactin complex at points of contacts in hippocampal neurons. J Neurochem 117:48–60.

38. Yamada H, Abe T, Satoh A, Okazaki N, Tago S, Kobayashi K, Yoshida Y, Oda Y, Watanabe M, Tomizawa K, Matsui H, Takei K. 2013. Stabilization of actin bundles by a dynamin 1/cortactin ring complex is necessary for growth of filopodia. J Neurosci 33:4514–4526.

39. Carman PJ, Dominguez R. 2018. BAR domain proteins — a linkage between cellular membranes, signaling pathways, and the actin cytoskeleton. Biophys, Rev 10:1587–1606.

40. Daumke O, Roux A, Haucke V. 2014. BAR domain scaffolds in Dynamin-mediated membrane fission. Cell 156:882–892.

41. Dowd GC, Mortuza R, Bhalla M, Van Ngo H, Li Y, Rigano LA, Ireton K. 2020. *Listeria monocytogenes* exploits host exocytosis to promote cell-to-cell spread. Proc Natl Acad Sci USA 117:3789–3796.

42. Gianfelice A, Le PH, Rigano LA, Saila S, Dowd GC, McDivitt T, Bhattacharya N, Hong W, Stagg SM, Ireton K. 2015. Host endoplasmic reticulum COPII proteins control cell-to-cell spread of the bacterial pathogen *Listeria monocytogenes*. Cell Microbiol 17:876–892.

43. Rigano LA, Dowd GC, Wang Y, Ireton K. 2014. *Listeria monocytogenes* antagonizes the human GTPase Cdc42 to promote bacterial spread. Cell Microbiol Epub ahead of print.

44. Theriot JA, Mitchison TJ, Tilney LG, Portnoy DA. 1992. The rate of actin-based motility of intracellular *Listeria monocytogenes* equals the rate of actin polymerization. Nature 357:257–260.

45. Damke H, Baba T, Warnock DE, Schmid S. 1994. Induction of mutant Dynamin specifically blocks endocytic coated vesicle formation. J Cell Biol 127:915–934.

46. Macia E, Erlich M, Massol RH, Boucrot E, Brunner C, Kirchhausen T. 2006. Dynasore, a cell-permeable inhibitor of dynamin. Dev Cell 10:839–850.

47. Herath TUB, Roy A, Gianfelice A, Ireton K. 2021. *Shigella flexneri* subverts host polarized exocytosis to enhance cell-to-cell spread. Mol Microbiol 116:1328–1346.

48. Dhanda AS, Yu C, Lulic KT, Vogl AW, Rauch V, Yang D, Nichols BJ, Kim SH, Polo S, Hansen CG, Guttman JA. 2020. *Listeria monocytogenes* exploits host caveolin for cell-to-cell spreading. mBio 11: pii: e02857–02819.

49. Henmi Y, Tanabe K, Takei K. 2011. Disruption of microtubule network rescues aberrant acti comets in dynamin 2-depleted cells. PLoS ONE 6:e28603.

50. Lee E, De Camilli P. 2002. Dynamin at actin tails. Proc Natl Acad Sci USA 99:161–166.

51. Verma S, Han SP, Michael M, Gomez GA, Yang Z, Teasdale RD, Ratheesh A, Kovacs EM, Ali RG, Yap AS. 2012. A WAVE2-Arp2/3 actin nucleator apparatus supports junctional tension at the epithelial zonula adherens. Mol Biol Cell 23:4601–4610.

52. Narumiya S, Thumkeo D. 2018. Rho signaling research: history, current status and future directions. FEBS Lett 592:1763–1776.

53. Peterson MD, Mooseker MS. 1992. Characterization of the enterocyte-like brush border cytoskeleton. J Cell Sci 102:581–600.

54. Fukumatsu M, Ogawa M, F. A, Suzuki M, Nakayama K, Shimizu S, Kim M, Mimuro H, Sasakawa C. 2012. *Shigella* targets epithelial tricellular junctions and uses a noncanonical clathrin-dependent endocytic pathway to spread between cells. Cell Host Microbe 11:325–336.

55. McNiven MA, Kim LC, Krueger WE, Orth JD, Cao H, Wong TW. 2000. Regulated interactions between dynamin and the actin-binding protein cortactin modulate cell shape. J Cell Biol 151:187–198.

56. Mooren OL, Kotova TI, Moore AJ, Schafer DA. 2009. Dynamin 2 GTPase and cortactin remodel actin filaments. J Biol Chem 284:23995–24005.

57. Lamason RL, Welch MD. 2017. Actin-based motility and cell-to-cell spread of bacterial pathogens. Curr Opin Microbiol 35:48–57.

58. Belotserkovsky I, Sansonetti PJ. 2018. *Shigella* and Enteroinvasive Escherichia Coli. Curr Top Microbiol Immunol 416:1–26.

59. Walker DH, Ismail N. 2008. Emerging and re-emerging rickettsioses: endothelial cell infection and early disease events. Nat Rev Microbiol 6:375–386.

60. Duncan-Lowey JK, Wiscovitch AL, Wood TE, Goldberg MB, Russo BC. 2020. Shigella flexneri disruption of cellular tension promotes intercellular spread Cell Rep 33:108409.

61. Lamason RL, Bastounis E, Kafai NM, Serrano R, Del Alamo JC, Theriot JA, Welch MD. 2016. *Rickettsia* Sca4 reduces vinculin-mediated intercellular tension to promote spread. Cell 167:670–683.e610.

62. Engelbrecht F, Chun S-K, Ochs C, Hess J, Lottspeich F, Goebel W, Sokolovic Z. 1996. A new PrfA-regulated gene of *Listeria monocytogenes* encoding a small, secred protein which belongs to the family of internalins. Mol Microbiol 21:823–837.

63. Basar T, Shen Y, Ireton K. 2005. Redundant roles for Met docking site tyrosines and the Gab1 pleckstrin homology domain in InlB-mediated entry of *Listeria monocytogenes*. Infection and Immunity 73:2061–2074.

64. Ireton K, Payrastre B, Cossart P. 1999. The *Listeria monocytogenes* protein InlB is an agonist of mammalian phosphoinositude-3-kinase. J Biol Chem 274:17025–17032.

65. Dokainish H, Gavicherla B, Shen Y, Ireton K. 2007. The carboxyl-terminal SH3 domain of the mammalian adaptor CrkII promotes internalization of *Listeria monocytogenes* through activation of host phosphoinositide 3-kinase. Cell Microbiol 10:2497–2516.

66. Shen Y, Naujokas M, Park M, Ireton K. 2000. InlB-dependent internalization of *Listeria* is mediated by the Met receptor tyrosine kinase. Cell 103:501–510.

67. Kovacs EM, Makar RS, Gertler FB. 2006. Tuba stimulates intracellular N-WASP-dependent actin assembly. J Cell Sci 119:2715–2726.

68. Yao Q, Chen J, Cao H, Orth JD, McCaffery JM, Radu-Virgil S, McNiven MA. 2005. Caveolin-1 interacts directly with Dynamin-2. J Mol Biol 348:491–501.

